# A genome-wide CRISPR screen reveals how diatoms thrive in dynamic light

**DOI:** 10.1101/2025.09.15.676419

**Authors:** Jon Doenier, Dimitri Tolleter, Sarah Frail, Giovanni Finazzi, Adrien Burlacot, Ellen Yeh

**Affiliations:** Department of Biochemistry, Stanford School of Medicine, Stanford, CA 94305, USA; Biosphere Science and Engineering Division, Department of Plant Biology, Carnegie Science, Stanford, CA 94305, USA; Laboratoire de Physiologie Cellulaire et Végétale, Université Grenoble-Alpes, CNRS, CEA, INRAE, IRIG-DBSCI, 17 rue des Martyrs, 38000 Grenoble, France; Department of Biology, Stanford University, Stanford, CA 94305, USA; Department of Pathology, Stanford School of Medicine, Stanford, CA 94305, USA; Department of Microbiology & Immunology, Stanford School of Medicine, Stanford, CA 94305, USA; Chan Zuckerberg Biohub – San Francisco, San Francisco, California 94158, USA

## Abstract

Diatoms are a highly diverse group of phytoplankton that have a large impact on global primary production and carbon sequestration in the ocean^1,2^. However, they are evolutionarily divergent from model phototrophs of the green lineage, and limited screening tools have hampered discovery of unique diatom biology. To address this challenge, we developed a genome-wide CRISPR/Cas9 screen in the model marine diatom, *Phaeodactylum tricornutum.* The screen was applied to identify genes required for survival in different light regimes, including both high light and fluctuating light. We identified a broad set of uncharacterized genes, providing the foundation for mechanistic studies of diatom adaptation to dynamic light. Among these genes, we demonstrated that the red lineage-exclusive gene STROBE1 is a new potentiator of cyclic electron flow (CEF) required for CEF to generate a trans-thylakoid proton gradient^3,4^. As dynamic light conditions are common in marine environments, STROBE1 and other genes identified in this screen may contribute to the broad ecological success of diatoms^5,6^. This genome-wide genetic screen in *P. tricornutum* will accelerate the unbiased discovery of novel gene functions in these ecologically important organisms.

## Main

Oceans and other aquatic ecosystems are dynamic environments subject to surface winds, coastal upwellings, water column mixing, and plankton blooms. Under these conditions, light intensity can rapidly fluctuate between insufficient (limiting photosynthesis) and excessive (potentially causing oxidative damage). Among phototrophs, diatoms are particularly abundant in well-mixed, nutrient-rich marine environments such as coastal, upwelling, and polar regions^1,2^. Their ability to thrive in dynamic light environments^5,6^ suggests that diatoms have evolved unique molecular mechanisms to adapt to rapidly changing light.

During photosynthesis, light is used by photosystem (PS) I and II to generate linear electron flow (LEF) that produces NADPH and pumps protons into the thylakoid lumen. The light-induced proton gradient is, in turn, used to power ATP synthesis. However, NADPH and ATP production must be dynamically tuned to match energy needs for downstream carbon fixation reactions. Photosynthetic organisms have evolved many mechanisms to balance light-dependent energy production and carbon fixation. A common strategy in phototrophs is non-photochemical quenching (NPQ), in which excessive photons are dissipated as heat rather than used by the photosystems. NPQ is induced by the build-up of protons in the thylakoid lumen, that results in lumenal acidification. Electrons can also be redirected to alternative electron flows (AEF), which include cyclic (CEF), pseudo-cyclic (PCEF), and chloroplast-to-mitochondrial (CMEF) electron flows. By re-routing electrons away from LEF, these AEFs can regulate the rate of LEF, the ratio of ATP:NADPH produced, and the induction of NPQ.

While NPQ in diatoms is similar to that in model green phototrophs^7,8^, the molecular players and functions of AEFs in diatoms are different and largely unexplored^4^. AEFs are particularly suited to the timescale of response required for dynamic light (sec-min). In green phototrophs, CEF, PCEF, and CMEF have all been shown to contribute to energy homeostasis in dynamic light^9,10^. Diatoms are missing the genes required for PCEF and NADH dehydrogenase-like complex (NDH)-dependent CEF^4,11^. The key CEF regulators, proton gradient regulation (PGR) 5 and PGR-Like (PGRL) 1, are encoded in diatoms^6,12,13^. However, the very low CEF capacity in diatoms has raised doubts about its importance for acclimation to light^14^. CMEF has been proposed to have a constitutive role in balancing ATP and NADPH production in diatoms^14^, however its role in adaptation to dynamic light has not been explored so far. Finally, photosynthetic control relies on the lumenal pH to balance light-dependent energy production and carbon fixation^15,16^. Given its central role, diatoms may utilize new mechanisms to tune lumenal pH in dynamic light environments. Despite their ecological importance, gene function discovery in diatoms has been challenging due to a lack of unbiased screening tools. To discover evolutionarily divergent pathways in diatoms, we developed a genome-wide CRISPR/Cas9 screen in *P. tricornutum* and used it to identify genes required for survival in dynamic light.

## Results

### A genome-wide CRISPR/Cas9 screen in *P. tricornutum* enables gene function discovery

Cas9-based genome editing in *P. tricornutum* has been performed successfully by many research groups, however the efficiency was too low for a genome-wide screen^19,20^. We made improvements to Cas9 expression and sgRNA design to increase gene editing efficiency (Fig. 1a). First, we developed a transgenic *P. tricornutum* strain that constitutively expressed GFP-tagged Cas9. Selection for Cas9 was achieved by expressing Cas9-GFP and a bleomycin resistance gene from a single promoter using a P2A skip peptide to produce discrete protein products (Extended Data Fig. 1a)^17^. A non-targeting “decoy” sgRNA was also expressed to prevent non-specific genome editing by constitutively expressed Cas9, which was previously observed in other organisms^18^. Second, we identified the start site of the native *P. tricornutum* U6 promoter (+1) using 5’-RACE and showed that it is different than the +3 site previously used for sgRNA expression in *P. tricornutum*^19,20^ (Extended Data Fig. 1b). To compare the gene editing efficiency using different U6 start sites, we used sgRNAs against *ZEAXANTHIN EPOXIDASE 1* (*ZEP1*) involved in fucoxanthin biosynthesis^21^. The loss of brown fucoxanthin pigment in Pt *zep1* results in green colonies that can easily be scored: unedited wildtype (brown), partial disruption (mixed green and brown), or complete disruption (green) (Extended Data Fig. 1c). sgRNA expression using the native U6 start site (+1) drastically increased gene editing efficiency, resulting in ∼80% of colonies with partial or complete disruption of *ZEP1* compared to <5% using the previously annotated +3 site (Fig. 1b and Extended Data Fig. 1d). These modifications substantially improved the efficiency of *P. tricornutum* gene editing, enabling a genome-wide screen.

**Figure 1.**
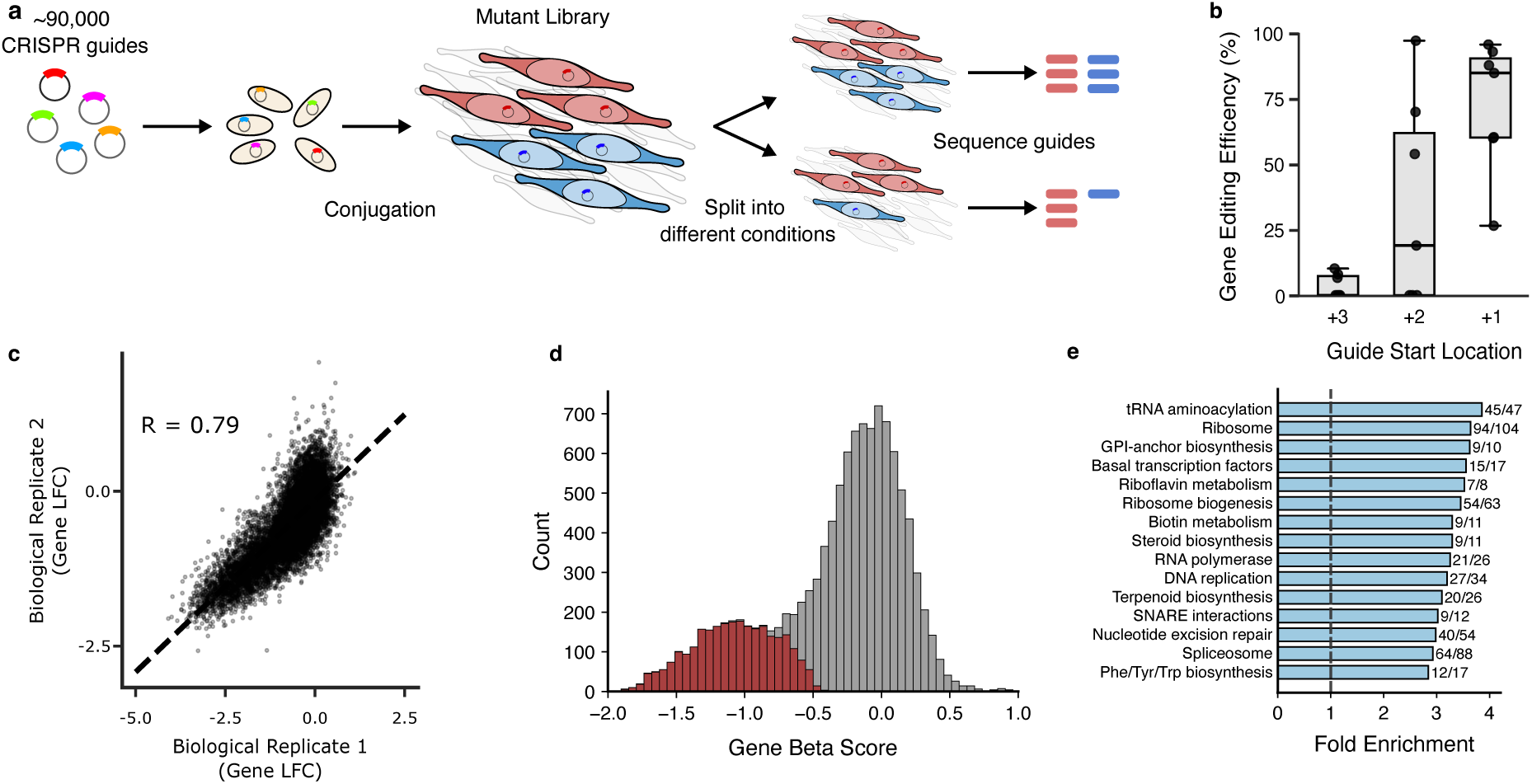
A genome-scale CRISPR/Cas9 screen in *P. tricornutum*. **a**, Overview of screen methodology. A library of sgRNA targeting the *P. tricornutum* genome are transformed into *E. coli* and then subsequently transferred into *P. tricornutum* using bacterial conjugation. The library of *P. tricornutum* mutants are grown in a single culture (red and blue represent two different mutants in the culture). The pooled mutant library is split into different growth conditions. After selection is complete, sgRNAs are sequenced to determine their abundance in different conditions. **b**, Efficiency of *zep1* disruption for guides using 3 different start sites: previously used start site (+3), alternative start site (+2), or native U6 promoter start site (+1). **c**, Comparison of gene depletion (log_2_-fold change) between two biological replicates of *P. tricornutum* mutant library grown under low light conditions. **d**, Histogram of estimated gene essentiality (beta-score) for *P. tricornutum* genes under low light. Red bars indicate genes with decreased growth rate that were statistically significant (q<0.01), while gray bars indicate genes that did not show significant decrease in growth rate. **e**, Function enrichment analysis of essential gene set showing top 15 pathways. For each pathway, the number of genes identified as essential versus the total number of genes with the functional annotation are shown.

We designed 76,057 sgRNAs targeting 11,252 protein-coding genes in the *P. tricornutum* genome (∼92% of all protein coding genes), resulting in an average of ∼6.8 guides per gene (Fig. 1a, Extended Data Fig. 1e-h, and Supplementary Table 1). 517 genes encoding repeat proteins with no unique targeting sites and 384 genes contained completely within another gene were excluded from the library (Extended Data Fig. 1g). The sgRNA library also included 5,000 non-targeting sgRNAs, 5,000 safe-targeting sgRNAs, and 4,148 essential gene targeting sgRNAs (Extended Data Fig. 1e and Supplementary Table 1). The library of sgRNAs was cloned into a plasmid library and transformed into *E. coli*, which were then conjugated to Cas9-expressing *P. tricornutum*. The resulting mutant library was split into different growth conditions and grown for approximately 12 generations (Fig. 1a). Biological and technical replicates showed a high correlation, demonstrating that the screen results are reproducible (Fig. 1c and Extended Data Fig. 2).

Under constant low light conditions, 2,727 genes (24.2% of targeted genes) caused a significant growth defect when mutated (Fig. 1d). This gene set was substantially enriched for essential functions, including DNA replication, RNA transcription, ribosomal translation, ribosome biogenesis, and aminoacyl-tRNA biosynthesis (Fig. 1e, Extended Data Fig. 3a, and Supplementary Table 2). Most *P. tricornutum* essential genes are conserved across eukaryotic taxa; however, a large number are conserved in diatoms and not shared with green phototrophs or other eukaryotes (Extended Data Fig. 3b). Identification of the set of *P. tricornutum* essential genes demonstrates the utility of the screen for biological discovery.

### CMEF importance is increased in high light

To investigate the molecular mechanisms that diatoms use to cope with dynamic light, we grew *P. tricornutum* mutant libraries under different light conditions: constant low (50 µmol photons m^-2^ s^-1^), medium (170 µmol photons m^-2^ s^-1^), or high (800 µmol photons m^-2^ s^-1^) light and fluctuating light (800 µmol photons m^-2^ s^-1^ for 1 minute, 20 µmol photons m^-2^ s^-1^ for 5 minutes). As high light is a component of the fluctuating light condition, we first sought to identify genes required for survival in constant high light. We identified 170 genes depleted under high light compared to the low light condition (Fig. 2a, Supplementary Table 3, and Supplementary Table 4). Many of the top genes point to adaptations in photosynthetic function that have been previously identified as key players for high light acclimation, including biogenesis and repair of photosystems (*FILAMENTOUS TEMPERATURE-SENSITIVE H*^22^, chloroplast (Cp) *FILAMENTOUS TEMPERATURE-SENSITIVE Y*^23,24^, Cp *SIGNAL RECOGNITION PARTICLE 54*^23,24^, *HYPERSENSITIVE TO HIGH LIGHT 1*^25^, *RESISTANCE TO PHYTOPHTHORA 1*^26^), carotenoid regulation and biosynthesis^27,28^ (*VIOLAXANTHIN DE-EPOXIDASE*, *CAROTENOID ISOMERASE 4*, *CAROTENOID ISOMERASE 5*), light-induced transcriptional regulation (Pt*AUREOCHROME* (*AUREO*) *1c*)^29^, antioxidant cofactors biosynthesis (*VITAMIN E 1*^30^, *GAMMA-TOCOPHEROL METHYLTRANSFERASE*^30^, *PYRIDOXINE BIOSYNTHESIS 1*^31^), and carbon concentrating mechanism^32^ (Pt*SOLUTECARRIER* (*SLC*) *4-5*, Pt*SLC4-6*, *PtCARBONIC ANHYDRASE* (*CA*) *2*, γ-*CA*) (Fig. 2a and Supplementary Table 4).

**Figure 2.**
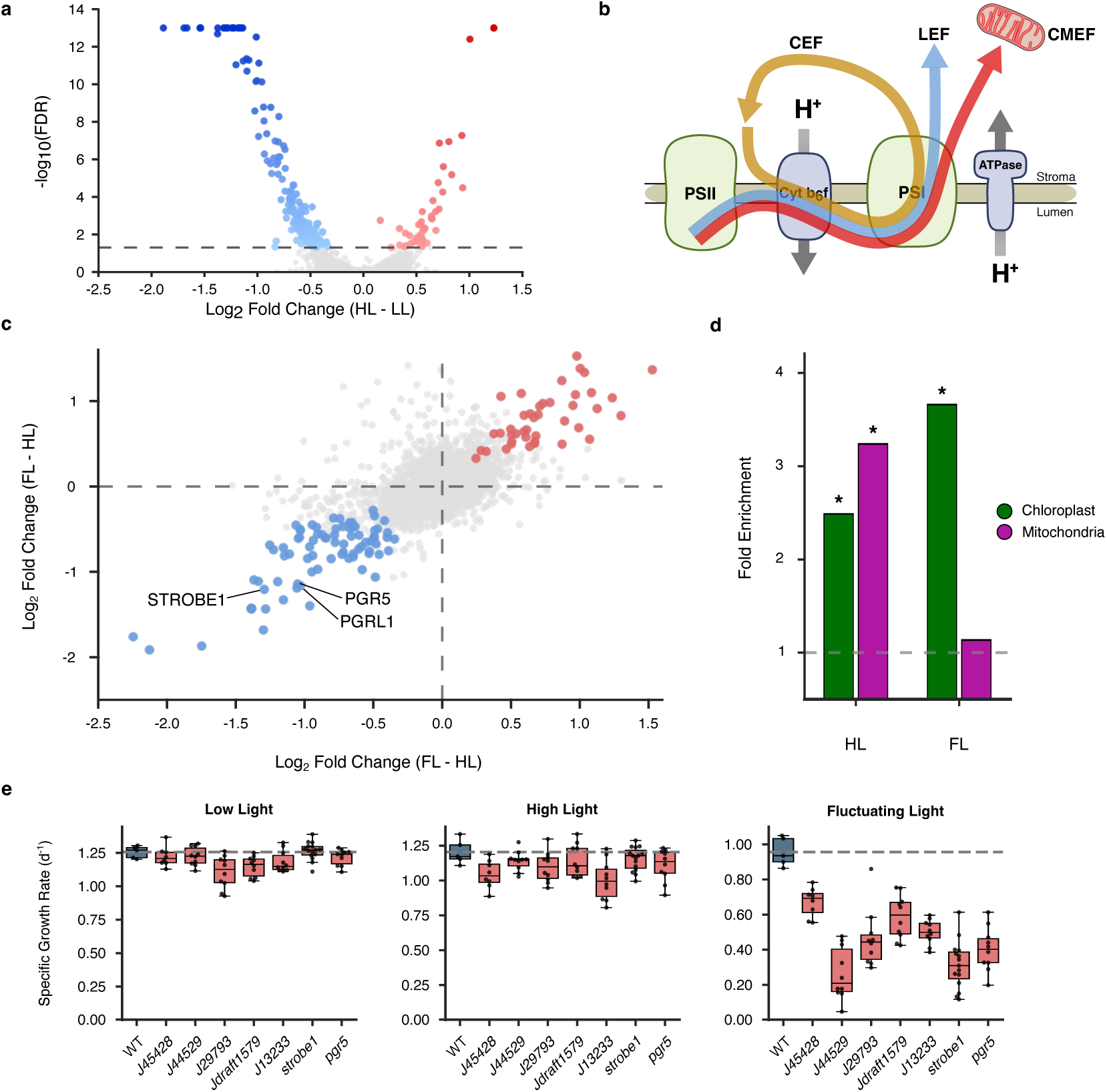
Screens for high light and dynamic light identify known and new genes. **a**, Volcano plot of high light compared to low light conditions showing genes depleted (blue), enriched (red), or unchanged in abundance (gray) under high light (q<0.05). Results represent 3 biological replicates, each with 2 technical replicates. **b**, Diagram of photosynthetic electron flows present in diatoms: linear electron flow (LEF, blue), chloroplast-to-mitochondria electron flow (CMEF, red), and cyclic electron flow (CEF, yellow). LEF and CMEF use both photosystems (PS) II and I, while CEF uses only PSI. Electron flow is coupled to proton pumping into the thylakoid lumen by cytochrome b6f. The proton gradient generated is used to produce ATP by ATP synthase. **c**, Relative gene abundance in fluctuating light compared with constant low light (x-axis) or high light (y-axis) conditions. Genes were depleted in dynamic light conditions (blue), enriched in dynamic light conditions (red), or unchanged (gray) with q<0.05. **d**, Enrichment of predicted chloroplast (green) or mitochondria (purple) genes in high light (HL) or fluctuating light (FL) conditions compared to the entire screen gene set. * signifies *p*<1x10^-4^. **e**, Specific growth rate of select mutants in low, high, and fluctuating light conditions. Results shown are median (center line), interquartile range (IQR, box), and 1.5xIQR (whiskers) for ≥2 independent mutants and ≥3 biological replicates per strain. Dashed line represents mean from WT cells.

Besides these mainly plastid proteins, the high light-depleted gene set is also enriched for mitochondrial proteins, including 15 of 18 nuclear-encoded subunits of Complex I (Fig. 2d and Extended Data Fig. 4a), suggesting a potential role of the CMEF. Consistent with this, *P. tricornutum* showed greater growth inhibition when treated with Complex I inhibitors in high light compared to low light (Extended Data Fig. 4b). It was previously shown that CMEF, in which photosynthetically-derived electrons are fed into the mitochondrial electron transport chain, has an important role in generating additional ATP without NADPH to optimize energy needs for carbon fixation (Fig. 2b)^14^. Our results show an increased requirement for CMEF in high light, suggesting that CMEF is the major AEF optimizing photosynthesis in diatoms mainly in conditions of saturating light intensities.

### Unbiased screen identifies new genes for survival in dynamic light

In addition to high-light acclimation, fluctuating light requires mechanisms to rapidly regulate photosynthesis. We identified 79 genes that were depleted in the fluctuating light condition compared to all constant light conditions as dynamic light-specific genes (Fig. 2c, Supplementary Table 3, and Supplementary Table 5). Unlike high-light acclimation, the dynamic light-specific genes were not enriched for mitochondrial genes, indicating that CMEF does not contribute specifically to dynamic light adaptation (Fig. 2d). Instead, the CEF effectors *PGR5* and *PGRL1* were identified as dynamic light-specific genes^4^. Two genes involved in NPQ regulation were also identified: 1) *ZEP3*, required for NPQ relaxation after high light exposure important for acclimation to fluctuating light^33^ and 2) the transcription factor *AUREO1a*^29^ which has a likely function in NPQ relaxation after high light exposure. Our results show that CEF and NPQ relaxation are important specifically under dynamic light, likely by lowering the lumenal pH during low-to-high light transition and relaxing NPQ mechanisms during high-to-low light transition, respectively.

The dynamic light-specific gene set also implicates previously unrecognized mechanisms. Two genes homologous to characterized plant genes that catalyze post-translational modifications of Calvin-Benson-Bassham (CBB) enzymes were identified: *THIOREDOXIN F* catalyzes the thiol reduction of multiple CBB enzymes in response to light and is involved in photosynthetic induction in plants^34–36^. *Phatr3*_*J46871* encodes a SET-domain protein with homology to methyltransferases that catalyze methylation of the small and large subunits of Rubisco^37,38^. Until now, there has been no insight into the functional consequence of Rubisco methylation. The identification of these genes indicates that dynamic post-translational regulation of carbon fixation reactions may be required to coordinate with dynamic photosynthetic energy production.

Finally, the majority of genes depleted in dynamic light were of unknown function and not previously identified in green phototrophs. To validate these candidates, we generated mutant strains for 6 of these genes that were highly depleted in the screen (Extended Data Fig. 5) and tested their growth phenotype under fluctuating light. A *pgr5* mutant was also generated for comparison^12^. All mutants exhibited a decrease in growth rate compared to wildtype (WT) when grown under fluctuating light conditions compared to constant low or high light conditions (Fig. 2e), demonstrating that the screen successfully identified new genes of unknown function required for survival in dynamic light.

### STROBE1 is a CEF potentiator

Of the newly validated genes, the growth defect of *Phatr3_EG00039* (hereafter *STROBE1*) was among the most specific to the fluctuating light condition (Fig. 2e). To verify the specificity of the *strobe1* defect, we generated a complementation strain by expressing STROBE1-GFP in the *strobe1* mutant background. STROBE1-GFP co-localized with chlorophyll (Extended Data Fig. 6a), consistent with a plastid localization predicted by ASAfind^39^. The growth of *strobe1;STROBE1-GFP* was also restored to WT levels in fluctuating light (Fig. 3a and Extended Data Fig. 6b). To investigate whether *strobe1* is affected in its capacity for photosynthesis, we measured PSII yield in response to light. When grown under constant light, *strobe1* showed a similar PSII yield as compared to WT over a range of light intensities (Extended Data Fig 7a). However, when grown under fluctuating light for 2h, *strobe1* displayed a nearly 50% decrease in maximal PSII yield (Fv/Fm) compared to WT, which was partially restored in *strobe1;STROBE1-GFP* (Fig. 3b). These results indicate that STROBE1 has a critical role in protecting PSII from photoinhibition and maintaining photosynthetic capacity in dynamic light conditions.

**Figure 3.**
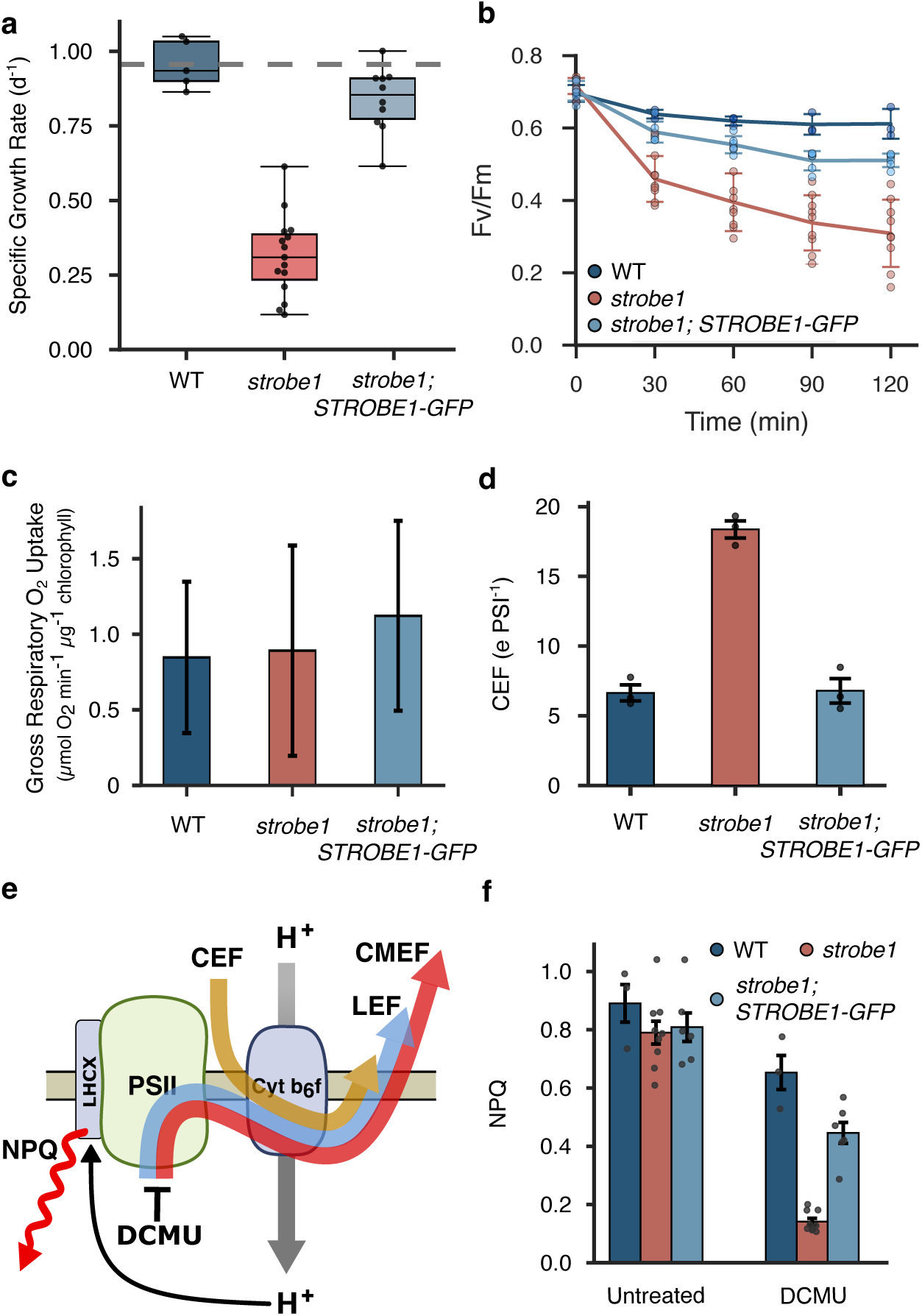
STROBE1 potentiates a CEF-dependent proton gradient. Comparison of wildtype (dark blue), *strobe1* (red), and *strobe1*;*STROBE-GFP* (light blue) for the following: **a**, Growth rate under fluctuating light. Results shown are median (center line), interquartile range (IQR, box), and 1.5xIQR (whiskers) for 5 biological replicates of ≥2 independent mutants. Dashed line represents mean from WT cells. **b**, Fv/Fm over 2 hours growth under fluctuating light. Results shown are mean +/-1 standard deviation for 3 biological replicates of ≥2 independent mutants. **c**, Difference of oxygen uptake in untreated cells (control) and cells treated with myxothiazol (myxo) and salicylhydroxamic acid (SHAM). Results shown are mean +/- 1 standard deviation for 4-8 biological replicates of 1 mutant strain. **d**, CEF activity in red-light adapted cells. Results shown as mean +/-1 standard deviation for 3 biological replicates of 1 mutant strain. **e**, Scheme showing NPQ induction by low lumenal pH. DCMU treatment blocks LEF (blue) and CMEF (red), while CEF (yellow) is active. **f,** NPQ after 5 minutes of high light treatment with no drug or with DCMU. Results shown are mean +/-1 standard deviation for 3 biological replicates of ≥2 independent mutants.

To investigate the STROBE1 mechanism, we measured its effect on the activity of LEF and AEFs. We first measured the capacity of *strobe1* to perform LEF and CMEF by measuring photosynthetic O_2_ exchange rates by membrane-inlet mass spectrometry (MIMS; Extended Data Fig. 9a-e). *strobe1* showed no defect in net O_2_ production compared to WT, indicating LEF was intact (Extended Data Fig. 9c). It also had a similar level of mitochondria-dependent O_2_ uptake as WT cells, ruling out STROBE1 function in CMEF (Fig. 3c and Extended Data Fig. 9a-e). These results are consistent with the screen results indicating a specific role for CMEF in high light adaptation. We next measured CEF activity by electrochromic shift (ECS)^40^, which showed 3-fold increased CEF activity in *strobe1* compared to WT (Fig. 3d and Extended Data Fig. 9f-h). High CEF capacity usually increases resilience to light stress, but surprisingly increased CEF activity in *strobe1* results in increased photoinhibition and reduced survival in dynamic light^16,41^.

CEF usually pumps protons into the lumen resulting in lumenal acidification, which powers ATP synthesis and induces NPQ. We assessed the lumenal pH in *strobe1* cells using NPQ as a sensor (Fig. 3e and Extended Data Fig. 7)^42^. As a control, we tested *pgr5* which showed reduced NPQ, a defect observed previously in *pgr5* in plants^43^ and *pgrl1* in green algae^44^ (Extended Data Fig. 7d,e)^45^. Interestingly, when all AEFs were active, no difference in NPQ was observed between WT, *strobe1*, or *strobe1;STROBE1-GFP* (Extended Data Fig. 7b,d). This result showed that NPQ is intact and can be induced upon dark-to-light transition in *strobe1* but also that the loss of STROBE1 may be compensated for by other AEFs (Supplementary discussion). We used the inhibitor 3-(3,4-dichlorophenyl)-1,1-dimethylurea (DCMU) to inhibit PSII-dependent electron transport, including LEF and CMEF, so that only CEF-dependent NPQ induction could be measured (Fig. 3e). When treated with DCMU, *strobe1* showed markedly reduced NPQ levels compared to WT or *strobe1;STROBE1-GFP*, suggesting a specific defect in the generation of low lumenal pH by CEF (Fig. 3f). Addition of the proton ionophore nigericin, which dissipates the proton gradient, abolished NPQ in all the conditions tested (Extended Data Fig. 8).

Altogether, *strobe1* does not show changes in LEF, CMEF, or NPQ. Rather, its effects on photosynthetic control are tightly linked with CEF. Increased CEF activity, without generation of low lumenal pH has not been observed in mutants of other AEF effectors. Hence, while STROBE1 activity is linked to CEF, it likely regulates lumenal acidification by CEF via a new mechanism. We propose that STROBE1 is a new CEF potentiator required for protons to accumulate in the lumen in a CEF-dependent manner.

### STROBE1 is restricted to red lineage phototrophs that are ubiquitous in modern oceans

We performed phylogenetic analysis of STROBE1 to explore its conservation across ecologically relevant phototrophs. Orthologs of STROBE1 were detected broadly in red lineage phototrophs of complex endosymbiotic origin represented by supergroups TSAR, Cryptista, and Haptista, including diatoms, dinoflagellates, haptophytes, and cryptomonads (Fig. 4a)^46^. However, it was not detected in genomes of rhodophytes that contain primary red plastids nor any green lineage phototrophs such as chlorophytes or land plants (Fig. 4a). Our analysis indicates that STROBE1 evolved in red lineage phototrophs after their divergence from the green lineage. In the red lineage, it either evolved from a red algal gene absent in representative rhodophytes or alternatively evolved from a gene acquired after endosymbiosis of an ancestral red alga, which then spread via additional endosymbiotic events^47^. A similar red lineage-specific pattern of evolution was observed for pyrenoid shell proteins (Pyshell), recently described structural proteins that enclose diatom pyrenoids and are required for specialized carbon concentration mechanisms^48,49^.

**Figure 4.**
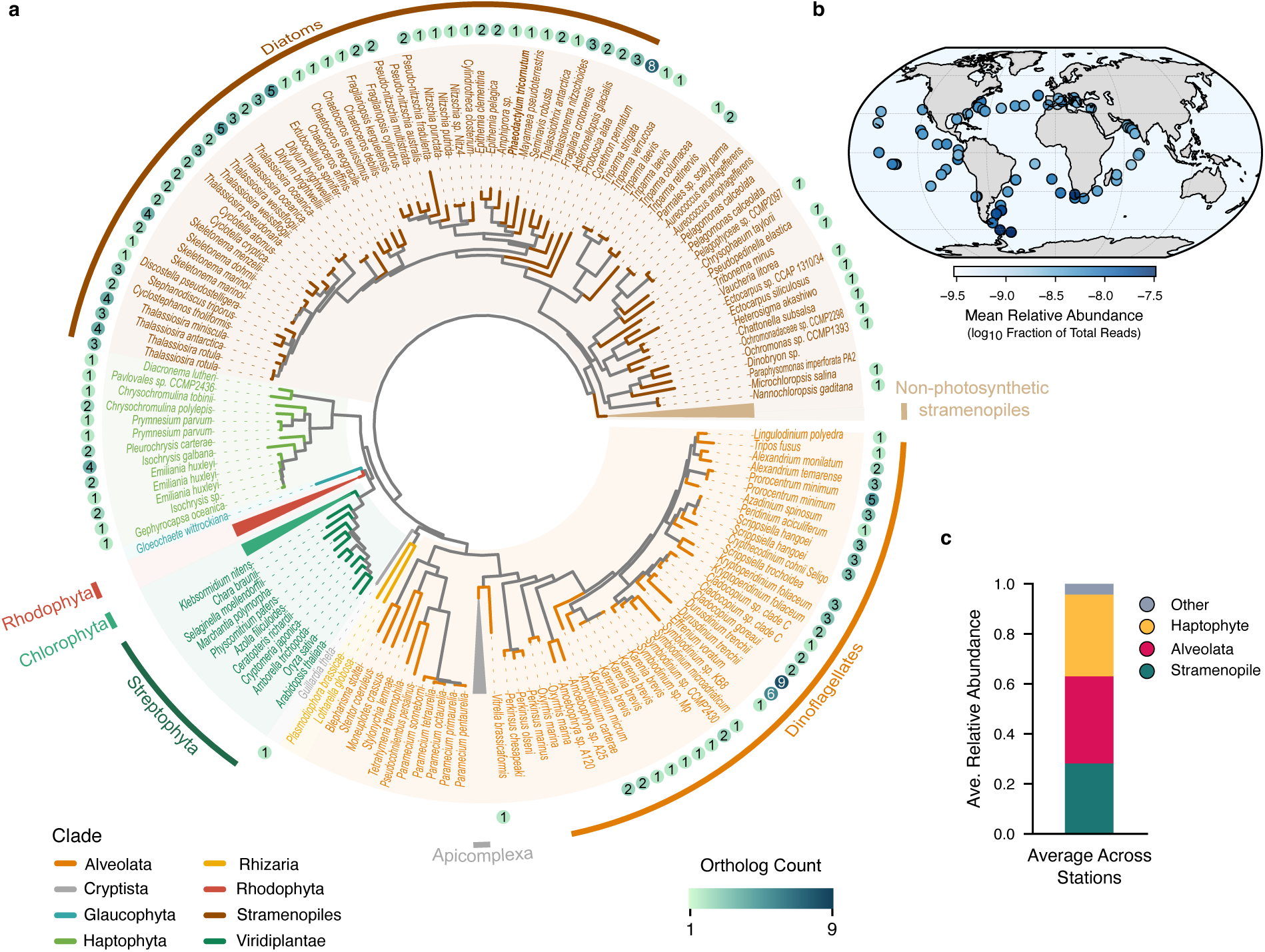
STROBE1 is globally expressed and restricted to red-lineage phototrophs. **a**, Phylogeny of STROBE1 orthologs across major eukaryotic clades. Clades are color-coded as indicated. Ortholog count per genome is shown in in teal-shaded circles on the outer ring. **b**, Global distribution of *in situ* STROBE1 expression across the Marine Atlas of Tara Oceans Unigene and meta-transcriptomic samples. Shading indicates log10-transformed relative abundance. **c**, Average taxonomic contributions to *STROBE1* transcripts across all stations.

The conservation of STROBE1 in eukaryotic phototrophs containing complex red plastids that dominate modern oceans, namely diatoms, dinoflagellates, and coccolithophores^1,50^, suggests it likely has an important function across ecologically important taxa. We investigated the expression of STROBE1 in metagenomic marine sampling data from the *Tara Oceans*. Expression of STROBE1 orthologs was detected at all marine sample sites (Fig 4b). Consistent with its phylogenetic distribution, *STROBE1* transcripts were mainly detected from groups containing red lineage algae: stramenophile, alveolata, and haptophytes (Fig 4c). This broad geographic expression indicates that STROBE1 is required ubiquitously in response to dynamic light.

## Discussion

### A milestone tool for exploring diatom biology

We developed a genome-wide CRISPR/Cas9 screen in the model diatom *P. tricornutum* and successfully performed a viability-based negative selection screen. Since negative selection screens have lower signal-to-noise characteristics and statistical power than positive selection screens, the screening protocol described herein meets a high bar of stringency for genome coverage, sensitivity, and reproducibility. We envision additional improvements to this first version of the screen that will further increase its efficiency, such as improving conjugation with alternative bacterial strains, integration of guides into the chromosome in safe-harbor locations, and improved sgRNA library design. Although genetically tractable, *P. tricornutum* is an atypical diatom lacking a silica cell wall in some morphotypes^51^. Hence, this genome-wide screen is best applied to identify traits that are conserved among diatoms. Future screens may identify biomarkers for ecological functions known to have specialized mechanisms in diatoms, such as nutrient acquisition^52–54^ or carbon concentration mechanisms^32,48,49^. Alternatively, screens for genes involved in secondary metabolite biosynthesis could create genetically-engineered algal strains to produce high-value lipids or carotenoids for biotechnology applications^55^. Overall, our work shows that the screen is robust and adaptable for the interrogation of diverse and new aspects of diatom biology.

### CMEF is critical in diatoms for high light acclimation

CMEF has been shown in diatoms to be the main pathway for ATP generation and regulation of the ATP:NADPH ratio to match the energy requirements of CO_2_ fixation^14^. Our findings show that CMEF is critical primarily to respond to constant high light intensities, conditions typical when diatoms reach an unmixed marine surface layer. While CMEF is required to respond to the high light component of fluctuating light, other pathways are required to cope with dynamic light specifically. Despite the importance of the CMEF pathway in diatoms, the molecules that shuttle electrons from the chloroplast to mitochondria and their associated transporters have yet to be identified^4,14^. Such a pathway in diatoms might involve multiple transporters to pass the four membranes of the chloroplast. The identification of a malate dehydrogenase (Phatr3_J42398), a 2-oxoglutarate transaminase (Phatr3_J8945), and glutamine synthetase (Phatr3_J22357) in the high-light-depleted gene set suggests that, like in plants, a malate shuttling might be involved in transferring reducing equivalents^56^. Though annotated malate transporters (Dit1/Phatr3_J8990, Phatr3_J23908, and Phatr3_EG02645) were not identified, several transporter genes were identified that may be candidates for a malate shuttle involved in CMEF (Phatr3_J11866, Phatr3_J43194, Phatr3_EG02269, Phatr3_J13994, and Phatr3_J47805). Our results open new opportunities for mechanistic studies of CMEF in diatoms.

### New CEF mechanisms in diatoms are key for survival in dynamic light

In green phototrophs, CEF, PCEF, and CMEF have all been shown to be key AEFs that rebalance the energy homeostasis in dynamic light environments^9,10,57^. The low CEF capacity in diatoms observed in previous studies appeared unlikely to be significant for regulating photosynthesis^14^. Here, we have identified that major CEF regulators such as PGRL1 and PGR5 were found in the dynamic light-specific gene set, strongly suggesting CEF is the major AEF required for survival of diatoms in dynamic light environments. Importantly, we identified STROBE1 as a new CEF potentiator required for survival in dynamic light that is exclusive to red lineage phototrophs.

How is low CEF capacity efficiently used for photoacclimation in diatoms? The unusual *strobe1* phenotype suggests that STROBE1 mediates an atypical mechanism not observed in green lineage phototrophs. Our results provide mechanistic models for further testing. First, STROBE1 is unlikely to be a new component of the PGR5-regulated CEF pathway. *strobe1* mutants showed increased CEF activity, which is unexpected if it participated in PGR5-regulated electron cycling around PSI. *pgr5* also does not phenocopy *strobe1* in its effect on NPQ: Loss of PGR5 disrupts NPQ in the absence of inhibitors when NPQ in *strobe1* is unaffected. Yet, *pgr5* showed a strong NPQ induction in the presence of DCMU when NPQ in *strobe1* is disrupted (Extended Data Fig. 7d). Our results are consistent with the presence of compensatory CEF activity under different conditions and suggests another CEF pathway, not dependent on PGR5, in *P. tricornutum*.

Next, we considered that STROBE1 might mediate this PGR5-independent CEF pathway, perhaps one similar to an NDH-dependent CEF which is missing in diatoms. In this model, the STROBE1-mediated CEF is more efficient than PGR5-regulated CEF, translocating more protons per electron. Its disruption leads to increased PGR5-regulated CEF in *strobe1* but is insufficient to compensate for the reduced proton pumping. Using photosynthetic electron flow measurements, we calculated the number of protons transported in *strobe1* compared to WT and estimated that the proposed STROBE1-mediated CEF would need to translocate at least 4.7 H^+^ per electron to account for the “missing” protons observed in *strobe1* mutants (Supplementary Discussion). NDH-dependent CEF has demonstrated the highest efficiency to date transporting 4 H^+^ per electron^58^. While it is possible that STROBE1 could mediate a new CEF pathway with greater efficiency than NDH-dependent CEF, additional components of a new pathway would need to be identified to support this model of STROBE1 function.

Finally, an alternative model for STROBE1 function is that it modulates CEF efficiency by controlling a leak of protons through the thylakoid membrane, effectively increasing H^+^ transported per electron by CEF (Fig. 5). Based on calculated electron and proton transport in the WT and *strobe1*, we estimate that STROBE1 may prevent the loss of 5-10% of transported protons across the thylakoid membrane (Supplementary Discussion), a magnitude consistent with the activity of uncoupling proteins that have been characterized^59^. Loss of STROBE1 would result in proton leakage out of the lumen and decrease the efficiency of CEF in generating lumenal acidification in dynamic light conditions (Fig. 5). Since a low lumenal pH slows down electron flow through the cytochrome b6f in a process called photosynthetic control, a higher lumenal pH would decrease photosynthetic control and stimulate CEF activity in *strobe1* mutants (Fig. 3)^15^. In dynamic light conditions, potentiation by STROBE1 of proton translocation efficiency would allow CEF to rapidly generate lumenal acidification. Hence, a relatively small amount of CEF can quickly limit photosynthesis and avoid the production of damaging photosynthetic electrons. Meanwhile, in constant high light, STROBE1 may uncouple the trans-thylakoid proton gradient to prevent the lumenal pH from becoming too low when CMEF is generating the cell’s ATP requirements.

**Figure 5.**
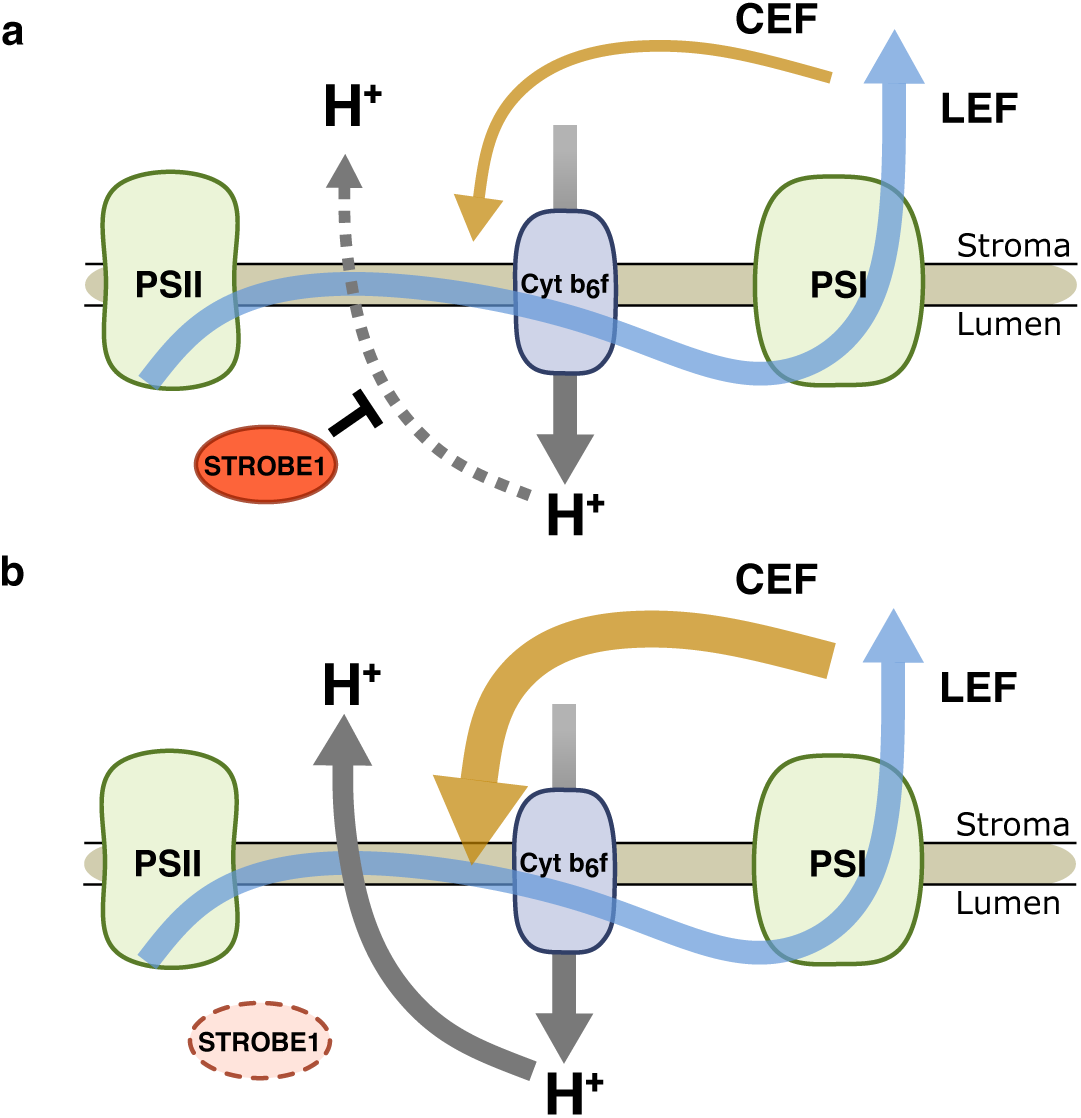
Proposed model of STROBE1 potentiation of CEF. **a**, STROBE1 is a negative regulator of proton translocation from the thylakoid lumen to the stroma (gray dashed arrow), facilitating generation of the trans-thylakoid proton gradient. **b**, Loss of STROBE1 increases the leak of protons into the thylakoid stroma (gray solid arrow), dissipating the proton gradient and causing a compensatory increase in CEF (yellow arrow).

Our work demonstrates the power of genome-wide screens in diatoms as a platform for uncovering core principles of photosynthetic adaptation to environmental conditions. Dynamic light dominates many marine ecosystems, the discovery of *STROBE1* and other diatom-specific genes may help explain the remarkable capacity of diatoms to outperform other phytoplankton groups in dynamic light conditions^60^. Further molecular characterizations of the genes reported here will reveal how aquatic photosynthesis is tuned to sustain planetary productivity.

## Methods

### Culturing

*Phaeodactylum tricornutum* (CCAP 1055/1) was grown at 18°C under full white LED lights (**Supplementary Fig. 1a**) with a 16 hour light, 8 hour dark diel cycle unless otherwise stated. Cultures were grown in sterile ESAW (Harrison, Waters, and Taylor 1980; Berges, Franklin, and Harrison 2001) modified to 0.88 mM NO3 and 40 µM PO4. Sodium metasilicate was omitted. Actively growing cells were used for all experiments.

### Conjugation

Conjugation was performed as published^61,62^ with minor modifications. To perform conjugation, actively growing liquid cultures of *P. tricornutum* were pelleted at 3,000 g for 10 minutes, resuspended in a small volume of ESAW, and plated on conjugation plates (1% agar (BD, Cat.# 214530), 5% Miller LB (BD, Cat.# 244620), ½ x ESAW, 0.88 mM nitrate, 0.48 mM ammonium). Plated *P. tricornutum* were grown for 1-2 days before conjugation. *E. coli* (Lucigen, Cat.# EC300110) harboring pTA-Mob^63^ and conjugation plasmids were cultured overnight in LB with appropriate antibiotics (20 µg/ml gentamycin (Goldbio, Cat.# G-400-1) and either 50 µg/ml kanamycin (Goldbio, Cat.# K-120-25) or 100 µg/ml carbenicillin (Sigma-Aldrich, Cat.# C1389-1G)). Bacterial cultures were diluted 1:50 in fresh antibiotic LB and grown to OD_600_ 0.8. The bacterial cultures were pelleted at 3,000 g for 5 minutes, resuspended in 37°C SOC media (NEB, Cat.# B9020S) and plated on top of *P. tricornutum* cultures. *P. tricornutum* and *E. coli* were mixed thoroughly and left until completely dry, approximately 10 minutes. Plates were incubated for 90 minutes in the dark at 30°C, then transferred to standard *P. tricornutum* growth conditions for 40-44 hours. *P. tricornutum* were harvested with ESAW and plated on selection media (conjugation plates supplemented with 50 µg ml^-1^ kanamycin or 25 µg ml^-1^ chloramphenicol (Sigma-Aldrich, Cat.# C0378) and 100 µg ml^-1^ Nourseothricin (Goldbio, Cat.# N-500-4) or 75 µg ml^-1^ Zeocin (Invivogen, Cat.# ant-zn-5)). Plates were sealed with parafilm and grown under standard conditions.

### Electroporation

Electroporation was performed as previously published^64^ with minor modifications. Electroporation plasmids were linearized with EcoRV-HF (NEB, Cat.# R3101S) and purified using Qiagen PCR cleanup kit (Qiagen, Cat.# 28506). Actively growing *P. tricornutum* (1-2x10^8^ cells) were pelleted (3000 g, 10 minutes) and washed 4 times with ice-cold 375 mM sorbitol. Cells were resuspended in 100 µL of ice-cold 375 mM sorbitol and mixed with ∼4 µg linearized electroporation plasmid and 40 µg boiled salmon sperm DNA (Rockland, Cat.# MB-103-1000). Cells were electroporated in a Gene Pulser XCell (BioRad) set to 7 square-wave pulses, 500 V, 5 ms pulse width, 1 s interval. Cells were recovered in 15 mL of ESAW and placed in the dark at 18 °C for ∼18 hours. Cells were then plated on appropriate selection plates, wrapped in parafilm, and grown under standard conditions.

### *P. tricornutum* Transgene Strain Generation

The Cas9-expressing strain was created by electroporating *P. tricornutum* with linearized pJD28.pUC19_diaCas9_GFP_2A_shBla as described above, and selecting on 75 µg mL⁻¹ Zeocin (Invivogen, Cat.# ant-zn-5). Cas9-GFP integration was confirmed by PCR and fluorescent microscopy. STROBE1 complementation strains were generated in the same way using linearized pJD135.pPtPBR11_NAT_FcpB_EF2_STROBE1_GFP and selected on 100 µg mL⁻¹ Nourseothricin (GoldBio, Cat.# N-500-4). Integration of the STROBE1-GFP cassette was confirmed by colony PCR and fluorescence microscopy.

### P. tricornutum Genome Editing

To generate single CRISPR/Cas9 genome edits, guides targeting the gene of interest were designed using Crispor^65^. Oligos (IDT) with 20 bp homology arms were amplified with prJD341.sgRNA_F and prJD137.sgRNA_R cloned into PtPuc3_diaCas9_sgRNA^19^ linearized with BsaI-HFv2 (NEB, R3733) using Gibson Assembly (NEB, Cat.# E2611L). Cloned plasmids were transformed into Stellar competent *E. coli* (Takara, Cat.# 636766). CRISPR KO plasmids were conjugated into *P. tricornutum* using the standard conjugation protocol. Colonies were screened for genome editing by PCR and sanger sequencing (McLab, South San Francisco, CA, USA). Heterozygous edits were confirmed by nanopore sequencing (Primordium Labs, Arcadia CA, USA). All CRISPR KO strains were regrown from a single cell to ensure a homogenous culture genotype.

### Plasmid Construction

Plasmid pJD28.pUC19_diaCas9_GFP_2A_shBla (GenBank Accession PV959482) was used for generation of the transgenic Cas9 expressing strain. To construct the plasmid, FcpB promoter and Cas9 were amplified from PtPuc3_diaCas9_sgRNA (Addgene, Cat.# 109219) and fused to GFP via PCR stitching. The shBle selectable marker was amplified from PtPuc3 (Addgene, Cat.# 62863). All fragments were assembled into a linearized pUC19 backbone using Gibson Assembly.

Plasmid pJD118.pRL1383_pt_NAT_U6 (GenBank Accession PV959483) was used as the cargo plasmid for conjugative delivery of the CRISPR/Cas9 sgRNA library. To construct the plasmid, a *P. tricornutum* centromere fragment was amplified from pPtPBR11 (Addgene, Cat.# 80386) and inserted into pRL1383a (Addgene, Cat.# 70692) using Gibson assembly. EF2 promoter and FcpC 5’UTR were amplified from genomic DNA. A NAT resistance gene was PCR-amplified from pNAT^66^, a FcpB terminator was amplified from pPtPBR11 and the sgRNA expression cassette was amplified from PtPuc3_diaCas9_sgRNA. All fragments were assembled into the centromere-containing pRL1383a backbone via Gibson Assembly. To enable subsequent cloning of the sgRNA library, a stuffer sequence containing blue-white screening elements was PCR-amplified from pUC19 and inserted between the U6 promoter and sgRNA scaffold via Gibson Assembly.

Plasmid pJD135.pPtPBR11_NAT_FcpB_EF2_STROBE1_GFP (GenBank Accession PV959484) was constructed as a rescue vector for complementation of STROBE1 mutants. The Nourseothricin resistance cassette from pNAT and an EF2-eGFP (Twist Bioscience, San Francisco, CA, USA) were PCR-amplified and Gibson-assembled into pPtPBR11. The STROBE1 coding sequence was amplified from genomic DNA and cloned in-frame between the EF2 promoter and GFP to yield the STROBE1–GFP fusion using NEBuilder HiFi DNA Assembly Master Mix (NEB, Cat.# E2621).

### U6 5’RACE

Total RNA from 30 mg of *P. tricornutum* was extracted (Qiagen Universal RNA Plus Kit, miRNA protocol, Cat.# 76404) and modified with a template switching reverse transcriptase (NEB, Cat.# M0466) using a template switching oligo (prJD302.TSO) and a converted to cDNA using a primer targeting the U6 gene (prJD306.U6_RT). cDNA was amplified (primers prJD303.TSO_amp, prJD307.U6_R), used as input for Zero Blunt II topo cloning (ABclonal, Cat.# RK30130), and transformed into Stellar competent *E. coli* (Takara, Cat.# 636766). Plasmid insert sequences were determined by Sanger sequencing (McLab, South San Francisco, CA, USA).

### DNA Extraction

*P. tricornutum* (25-250 × 10⁶ cells) were pelleted (3000 g, 10 minutes), resuspended in deionized water to a volume of 200 µL, and lysed with 200 µL lysis buffer (200 mM Tris-HCl pH 7.5, 50 mM NaCl, 25 mM EDTA, 1% SDS). Cells were disrupted with five freeze-thaw cycles using liquid nitrogen and a 65 °C heat bath, treated with 4 µL RNase A (20 mg mL^-1^, NEB, Cat.# T3018) for 15 minutes at 37 °C, and treated with 4 µL Proteinase K (20mg mL^-1^, Omega Bio-Tek, Cat.# C755C84) for 1 hour at 50 °C. DNA was extracted twice with phenol:chloroform:isoamyl alcohol (25:24:1; ThermoFisher, Cat.# AC327111000) and twice with chloroform (Sigma-Aldrich, Cat.# 366927), precipitated with 0.3 M sodium acetate, 0.7 vol isopropanol, 1 µL GlycoBlue (Invitrogen, Cat.# AM9515); pelleted (18,000 g, 30 min, 12 °C), washed 70% ethanol, air-dried 3 min, and dissolved in 10 mM Tris-HCl pH 8.0.

### Whole Genome Sequencing and SNV Identification

Genome-wide sequencing was performed on *P. tricornutum* CCAP 1055/1 to determine the locations of single-nucleotide polymorphisms (SNPs) present in the strain to inform sgRNA guide design. Genomic DNA sequencing libraries were prepared with NEBNext Ultra II FS DNA Library Prep Kit (NEB, Cat.# E6177S) using the manufacturer protocol for inputs >= 100ng. The library was sequenced on an Illumina MiSeq v3 flow cell (2 × 150 bp). Reads were trimmed with fastp v0.23.4^67^, aligned to the Phaeodactylum tricornutum ASM15095v2 reference using BWA-MEM v0.7.17^68^, and SNVs were called with FreeBayes v1.3.6^69^ and filtered using VCFtools v0.1.16^70^. Raw reads are available in the NCBI Sequence Read Archive (SRA) under accession SRR34985825.

### Genome-wide sgRNA guide design

sgRNA guides for the genome-wide CRISPR/Cas9 screen were designed using a custom pipeline. Candidate sgRNA sequences were identified via CRISPOR v4.99^65^ and further scored using CRISPRon V1.0 ^71^. A composite “design score” was computed based on the CRISPRon score penalized by predicted off-targets and position from the 3’end of the gene. Guides were checked by aligning their sequences against the genome using BWA v0.7.17 and were eliminated if there were off targets within 0 or 1 nucleotide changes, variants in the guide sequence, or if the guide overlapped with a higher scoring guide by more than 20%. The top 7 guides were kept for each gene. Positive-control guides targeting putative essential genes applied the same filters but did not enforce the overlap rule and kept all passing guides, whereas non-targeting controls were produced by shuffling protospacers until no genomic match remained and then randomizing the PAM’s first base. Safe-targeting guides were designed against regions of the genome that showed high H3K27 methylation^72,73^ and no RNA seq expression (Bio-sample accession numbers SAMN04488978-SAMN04489007 and SAMN06350641-SAMN06350652).

### Phaeodactylum tricornutum genome-wide screen

*E. coli* EPI300 Δ*sufA* harboring pTA-MOB and the sgRNA library was grown in LB Miller (BD, Cat.# 244620) supplemented with 50 µg mL⁻¹ kanamycin (GoldBio, Cat.# K-120-25) and 20 µg mL⁻¹ gentamicin (GoldBio, Cat.# G-400-1) at 37 °C, 250 rpm to an OD₆₀₀ of 0.9. Cells were pelleted (3,000 g, 5 min), washed twice in SOC medium (NEB, Cat.# B9020S) and resuspended in 37°C SOC. 4.4 L of *P. tricornutum* (4-7x10^6^ cells mL^-1^) were harvested by centrifugation (3,000 × g, 15 min), resuspended in 6 mL ESAW, and 250 µL aliquots were plated onto high-efficiency conjugation agar (1.4% BD Bacto agar, 30% salinity ESAW, 5% LB Miller, 0.88 mM nitrate, 0.48 mM ammonium). Plates were dried for 15 min, overlaid with the prepared E. coli suspension, incubated at 30 °C in the dark for 4 h, then shifted to 18 °C under 75 µmol photons m⁻² s⁻¹ for 40 h. *P. tricornutum* were harvested with ESAW and plated on selection media (conjugation plates supplemented with 50 µg ml^-1^ kanamycin (Goldbio, Cat.# K-120-25) and 125 µg ml^-1^ Nourseothricin (Goldbio, Cat.# N-500-4), and 75 µg ml^-1^ Zeocin (Invivogen, Cat.# ant-zn-5)). Plates were sealed with parafilm and grown under standard conditions for 2 weeks. Conjugates were scraped into ESAW and washed and pelleted (800 g, 5 min) three times to remove bacterial cells. Mutant libraries were recovered overnight in 500 mL ESAW supplemented with 50 mM Tris-HCl pH 8.2, 50 µg mL⁻¹ kanamycin and 125 µg mL⁻¹ nourseothricin (GoldBio, Cat.# N-500-4), then diluted to 3.6×10⁵ cells mL⁻¹ (∼2,000 x library coverage, ∼1.8×10⁸ total cells) and grown in technical duplicate at 18 °C with stir bar agitation (120 rpm) under four white led (**Supplementary Fig. 1b**) light regimes – low (50 µmol photons m⁻² s⁻¹), medium (170 µmol photons m⁻² s⁻¹), high (800 µmol photons m⁻² s⁻¹) and fluctuating (alternating 5 min at 20 µmol photons m⁻² s⁻¹ and 1 min at 800 µmol photons m⁻² s⁻¹) – maintaining cell density between 3.6×10⁵ and 1×10⁷ cells mL⁻¹ by periodic dilution. After ∼12 doublings, cultures were harvested (3,000 g, 10 min), pellets containing ≥1.8×10⁸ cells (∼2000 x library coverage) were flash-frozen, and DNA was extracted as above. For sequencing library preparation, episomal guides were first enriched by PCR with primers prJD59.pRL1383a_R and prJD459.Nat_F (20 cycles, Q5 Hot Start Polymerase, NEB, Cat.# M0494), followed by guide amplification to add Illumina trueseq adapters (prJD192–199, 12 cycles). PCR products were purified with a PCR cleanup kit (Qiagen, Cat.# 28506), indices were added using NEBNext Ultra II Q5 Master Mix (6 cycles; NEB, Cat.# E7805S/L), and final libraries were cleaned with AMPure XP beads (Beckman Coulter, Cat.# A63880). Libraries were sequenced on an Illumina NovaSeq X system using an 8-lane 10B flow cell (1 x 100bp). Data are available under BioProject PRJNA1305626.

### Genome-wide screen analysis

Adapter and quality trimming were performed with fastp v0.23.4 ^67^, and sgRNA read counts were obtained using MAGeCK count v0.5.9 ^74^ with median normalization. Log_2_ fold-changes relative to the untransformed plasmid library were calculated for each sample, and gene-level fitness effects (β-scores) were estimated by MAGeCK MLE v0.5.9 ^75^ using default parameters and the untransformed plasmid library as reference. Genes exhibiting negative β-scores with a Benjamini–Hochberg–adjusted FDR ≤ 1% were designated as essential. To identify condition-specific phenotypes, β-scores were quantile-normalized across conditions and pairwise differences were computed. P-values were derived via the “quantile matching” σ-estimation protocol ^76^ and adjusted by the Benjamini-Hochberg procedure to a 5% FDR.

KEGG pathway analysis was performed on the essential-gene set (β < 0, FDR ≤ 1%). Kegg ontology identifiers were assigned to all *P. tricornutum* genes using the merged outputs from BlastKOALA v3.1 and GhostKOALA v3.1 ^77^. Enrichment of essential genes was evaluated with gseapy v0.12.0 ^78^ in Enrichr mode (one-tailed hypergeometric test) against a background of all genes quantified with >1 sgRNA. P-values were adjusted by the Benjamini-Hochberg method; pathways with FDR < 0.05 were deemed significantly enriched.

### Bacterial Genome Editing

Deletion of the *sufA* locus in *E. coli* EPI300 was performed via λ-Red recombineering using the pREDTAI system (Addgene, Cat.# 51627). A 1.2 kb chloramphenicol resistance cassette was PCR-amplified from pKD3 (Addgene, Cat.# 45604) with 50 bp homology arms flanking the *sufA* open reading frame using Q5 Hot Start Polymerase (NEB, Cat.# M0494). The amplicon was purified using Qiagen PCR cleanup kit (Qiagen, Cat.# 28506) and 100 ng was electroporated into mid-log pREDTAI-harboring cells induced with 0.1% arabinose at 30 °C for 60 min. Electrocompetent cells were prepared by washing twice in ice-cold 10% glycerol. Cells were electroporated at 1.8 kV, 25 µF, 200 Ω (Bio-Rad Gene Pulser XCell). Following recovery in SOC medium (NEB, Cat.# B9020S) at 37 °C for 1 h, transformants were selected on LB agar containing 25 µg mL⁻¹ chloramphenicol (Sigma-Aldrich, Cat.# C0378). To cure pREDTAI, colonies were grown at 42 °C for 1 h and plated on antibiotic-free LB to isolate marker-retaining, helper-plasmid–free clones. Successful deletion of *sufA* was verified by colony PCR using primers prJD317.SufA_genotype and prJD318.SufA_genotype flanking the deletion junction and confirmed by Sanger sequencing.

### Growth curves

Growth curves were performed by inoculating 30 mL of ESAW medium with actively growing *P. tricornutum* strains to an initial OD₇₂₀ of 0.01. Cultures were incubated at 18 °C with continuous shaking (120 rpm) under one of three white led (**Supplementary Fig. 1b**) illumination regimes: constant low light (50 µmol photons m⁻² s⁻¹), constant high light (800 µmol photons m⁻² s⁻¹), or fluctuating light (5 min at 20 µmol photons m⁻² s⁻¹ followed by 1 min at 800 µmol photons m⁻² s⁻¹). Culture OD₇₂₀ was measured daily until values exceeded 0.20. Specific growth rates (µ) were calculated from the slope of the natural log of the OD₇₂₀ measurements versus time. A minimum of three biologically replicates were performed for each strain, and at least two independently generated mutants were measured for each gene.

### Chlorophyll fluorescence

Chlorophyll fluorescence was measured using a pulsed-amplitude-modulation (PAM) fluorimeter (Dual-Klass NIR, Walz GmbH, Germany) with the red measuring head. Detection pulses (10 µmol photons m^-2^ s^-1^ blue light) were supplied to measure fluorescence (F) at a frequency of 20-100 Hz. Red saturating flashes (4000 µmol photons m^-2^ s^-1^, 300 ms, 620 nm) were delivered to measure FM (maximal fluorescence yield in the dark-acclimated samples) and FM′ (upon actinic light exposure). Fluorescence emission was detected using a long-pass filter (>700 nm). The operating yield of photosystem II (Y(II)) was calculated as (FM′-F)/FM′, and NPQ was calculated as (FM-FM’)/FM′. The cell concentration of the samples was between 2-3x10^6^ cells/ml. HCO_3-_was added immediately before all PAM measurements (5 mM final concentration). When used, DCMU (10 µM final concentration) was added 10 seconds before the start of the light phase, after the measurement of FM. When used, nigericin (100 µM final concentration) was added 5 minutes before PAM measurements. When used, propyl Galate (PG, 1 mM final concentration) was added 1 minute before PAM measurements. Cells were dark-adapted for ten minutes before starting measurements.

### Gas exchange measurements

Gas exchange rates were measured using MIMS. Cells grown in air were collected during the exponential phase by centrifugation at 450 g for 3 min and resuspended in 1.5 mL of fresh buffered minimal medium (pH 7.2) at 30 μg chlorophyll mL^−1^. HCO_3−_ was subsequently added to the cell suspension (10 mm final concentration). The cell suspension was then placed in the MIMS reaction vessel (mounted with a 1 mil Teflon membrane), 100 *µ*L of ^18^O-enriched O_2_ (97% ^18^O Sigma-Aldrich, ref 490474-1L) was bubbled in the suspension. After closing the vessel, gas exchange was recorded during a dark-to-light transition (1,300 µmol photons m^−2^ s^−1^; green LEDs picked at 525 nm; Luminus reference PT-121-G-L11-MPK). Light intensity was set to be saturating using weakly absorbed green light for better homogeneity. Calculations of gross and net photosynthesis were done using the MIMS analysis software as previously described^79^. In some experiments, salicylhydroxamic acid (SHAM, 400 µM final concentration), myxothiazol (Myxo, 2.5 µM final concentration), dibromothymoquinone (DBMIB, 10 µM final concentration), or propylgallate (PG, 1 mM final concentration), or DCMU (10 µM final concentration) were added.

### Electrochromic shift measurement

For electrochromic shift measurements (ECS), cells were maintained at 19 °C under a light intensity of 40 µmol photons m⁻² s⁻¹, with a 12-hour light/12-hour dark photoperiod and continuously shaken at 90 rpm. Actively growing cells were harvested and concentrated to a density of 10 - 12 × 10⁶ cells mL^-1^. ECS signals were measured using a JTS-10 spectrophotometer (Biologic, France), equipped with a white probing LED and the appropriate interference filters (3-8 nm bandwidths). Cells were first put in darkness and exposed to single-turnover saturating flashes (using a frequency-doubled Nd:YAG laser with dyes, 700 nm, <10NS and >30 mJ) to measure the ECS signal induced by transferring only one electron per photosystem (ECS_stsf_). Cells were subsequently exposed to red light (320 µmol photons m⁻² s⁻¹) for 5 minutes, after which light was switched off and ECS recorded. Absorption difference signals (ΔI/I) were measured at 554, 520, and 565 nm (noted as [554], [520], and [565], respectively). ECS signals were corrected for the cytochrome c signal^40^ as: ECS = [520] – 0.25 × cyt c, where cyt c = [554] - 0.4 × [520] - 0.4 × [565]^14^. Under continuous illumination, the ECS originates from membrane potential generation by photochemical reactions and its dissipation by the chloroplast ATP synthase. Upon turning off the light, photochemistry ceases immediately, while ATP synthase dissipates ECS; hence, the immediate rate of ECS change (J) is related to the number of electrons flowing through photosystems before light is turned off. The rate of electron flow per photosystem was calculated as 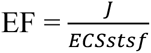. DCMU was added (10 µM final concentration) when calculating the rate of CEF.

### Phylogenetic analysis

The NCBI datasets command line tool v16.40.1^80^ was used to fetch proteomes of all annotated, non-MAG algal species in major clades (TaxIDs: SAR 2698737, Chlorophyta 3041, Haptista 2608109, Rhodophyta 2763, Cryptophyceae 3027), along with a limited list of 11 representative streptophytes. Translated transcriptome assemblies of other algal species were downloaded with iRODS iget v4.1.9 from the Marine Microbial Transcriptome Sequencing Project database^81^ hosted on CyVerse^82^. To check for completeness, these proteomes were analyzed for the presence of Eukaryotic and clade-specific conserved single-copy orthologues (--auto-lineage-euk) using BUSCO v5.3.2^83^. A species was removed from subsequent analysis if its proteome contained fewer than 50% Eukaryotic BUSCOs and fewer than 90% of clade-specific BUSCOs. As a final filtering step, the remaining proteomes were manually deduplicated to remove redundant assemblies representing the same or highly similar strain. This resulted in a final list of 351 proteomes used for species tree construction and orthologue inference.

The BUSCO phylogenomics pipeline was used to construct a species tree to identify single-copy complete Eukaryotic BUSCO proteins present in at least 75% of species and align, trim, and concatenate them into a supermatrix with associated partitions. IQ-TREE v2.3.6^84–86^ was run on this partitioned supermatrix with ModelFinder for 100 rounds of phylogenetic bootstraps.

To identify orthologues of the *P. tricornutum* STROBE1, Orthofinder v3.0.1b1^87^ was run with the core/assign feature using 54 representative species as the core set and assigning the remaining 297 species to the orthogroups identified in that core set. Orthogroups were manually investigated and one orthogroup composed of dinoflagellate genes was manually added to the STROBE1 orthogroup based on clearly conserved AlphaFold^88^ predicted tertiary structures. A full taxonomic breakdown of each species’ proteome was extracted from the NCBI Taxonomy database using the R tool taxize v0.10.0^89^. The R package ggtree was used to visualize the species tree and accompanying data^90^.

To identify global distribution of STROBE1, STROBE1 orthologs were aligned with MAFFT v7.525^91^ (--genafpair –maxiterate 1000), and the resulting alignment was converted to a profile HMM with hmmbuild v3.4. This model was uploaded to the Ocean Gene Atlas v2.0 and searched against the MATOU_v1_metaT dataset with an E-value cutoff of 1 x 10^-10^ and normalization set to “percent of total reads”^92^.

## Supporting information

Supplemental Discussion

## Acknowledgements

E.Y. is a Chan-Zuckerberg Biohub—San Francisco Investigator and supported by Burroughs Wellcome Fund. A.B. is supported by the Carnegie Institution for Science and DOE award DE-SC0019417. G.F. is supported by the European Research Council (ERC) advanced grant ‘‘ChloroMito’’ (grant number: 833184).

## Authors contribution

J.D., E.Y, and A.B. wrote the manuscript with input from all authors. J.D and E.Y. designed the CRISPR screen. J.D. and A.B. designed the light fluctuation screen and photosynthetic characterizations. Genome-wide screen, molecular biology, strain generation, and PAM were performed by J.D. MIMS was performed by D.T. ECS measurements were performed by D.T. and G.F. Phylogenomic analysis was performed by S.F. Photosynthetic measurements were supervised by A.B. All authors read and approved the final paper.

---

**Extended Data Figure 1.**
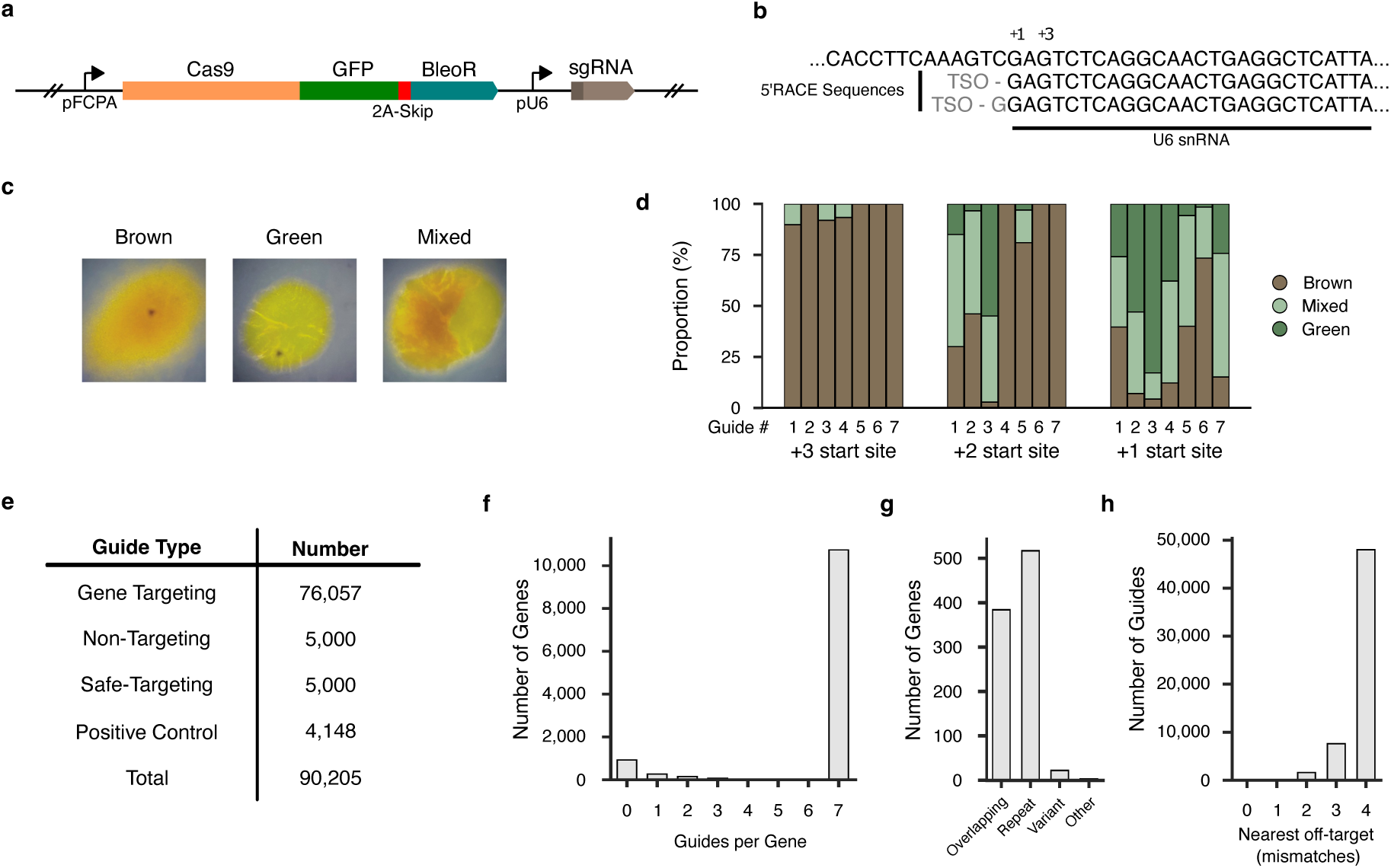
Development of a genome-wide CRISPR screen in *P. tricornutum*. **a**, Cas9 expression cassette consisted of the FcpA promoter driving expression of Cas9-GFP and bleomycin resistance gene separated by a P2A skip peptide and U6 promoter driving expression of a non-targeting sgRNA. **b**, Map of the Pt *U6 snRNA* gene locus showing the native start site (+1) and previously annotated start site for sgRNA expression (+3), top sequence. The native start site was mapped based on results of 5’ RACE: sequences following the template switching oligo (TSO; gray) were derived from the U6 snRNA (middle sequence) or non-templated G followed by U6 snRNA sequence (lower sequence). **c**, Classification of *zep1*-edited colonies: unedited colonies (left) appear brown when grown on agar plate. Complete *zep1* editing results in green colonies (middle). Partial *zep1* editing results in distinct regions of brown and green, which were scored as mixed (right). **d**, *zep1* editing efficiency for 7 sgRNAs expressed using different U6 promoter start sites. Colonies were classified as no gene editing (brown bars), complete gene editing (dark green bars), and partial gene editing (light green bars). Results represent >30 colonies per condition. **e**, Composition of sgRNA library targeting the *P. tricornutum* genome. Gene targeting guides are designed to target protein-coding genes. Non-targeting guides do not target any sequence in the genome. Safe-targeting guides target intergenic regions. Positive control guides target putatively essential genes. **f-h**, Characteristics of sgRNA library targeting the *P. tricornutum* genome: number of guides per gene (**f**). genes not targeted were either completely encoded within other genes (overlapping), had no unique coding sequence (repeat), or had a very high number of heterozygous SNVs (variant) (**g**). Distribution of sgRNAs based on the number of nucleotide differences compared to their most similar off-target sequence in the genome (**h**).

**Extended Data Figure 2.**
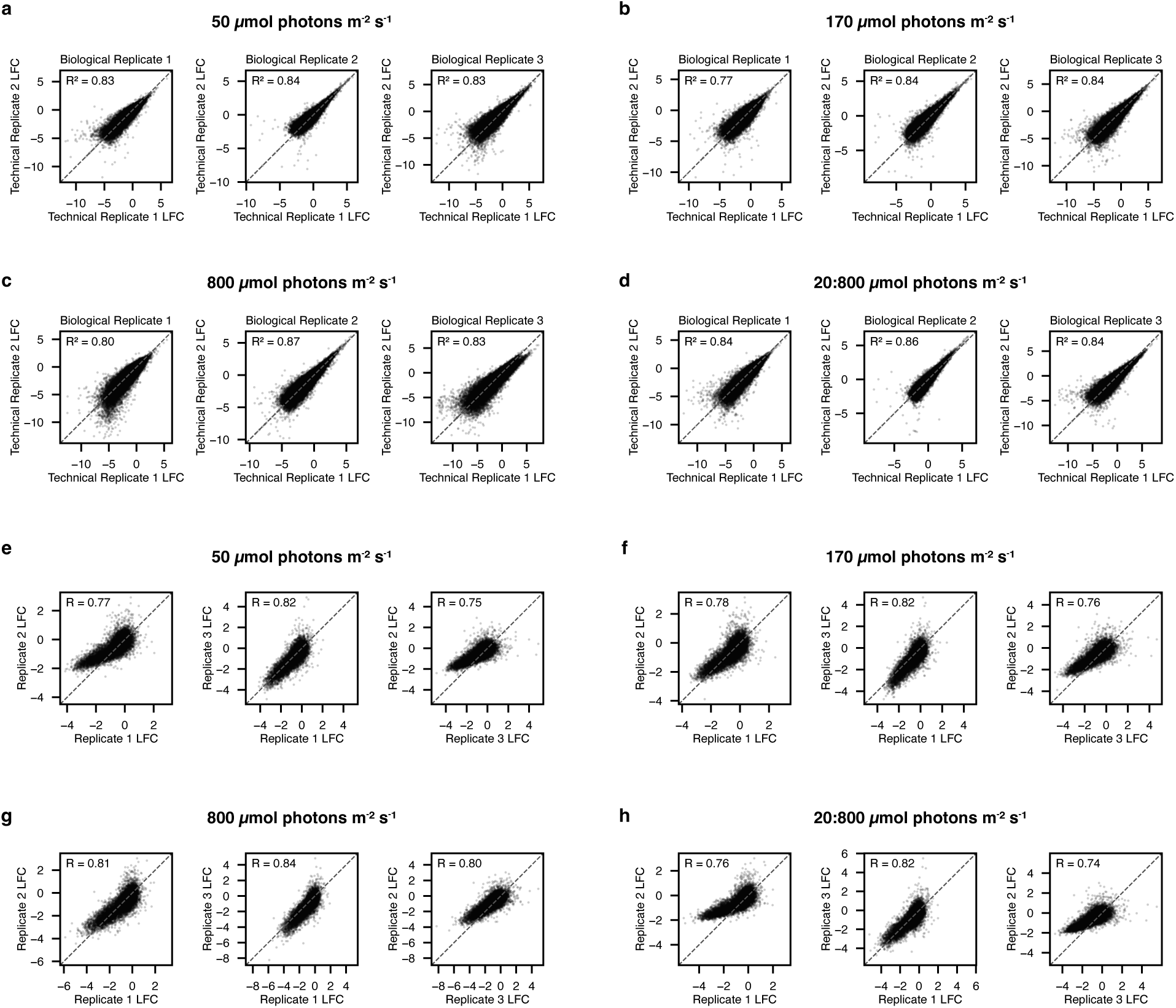
*P. tricornutum* genome-wide screen is highly reproducible. **a-d**, Comparison of log_2_ fold sgRNA abundance between technical replicates for low, medium, high, and fluctuating light conditions. **e-h**, Comparison of log_2_ fold gene summarized abundance (average over all sgRNA) between biological replicates for low, medium, high, and fluctuating light conditions.

**Extended Data Figure 3.**
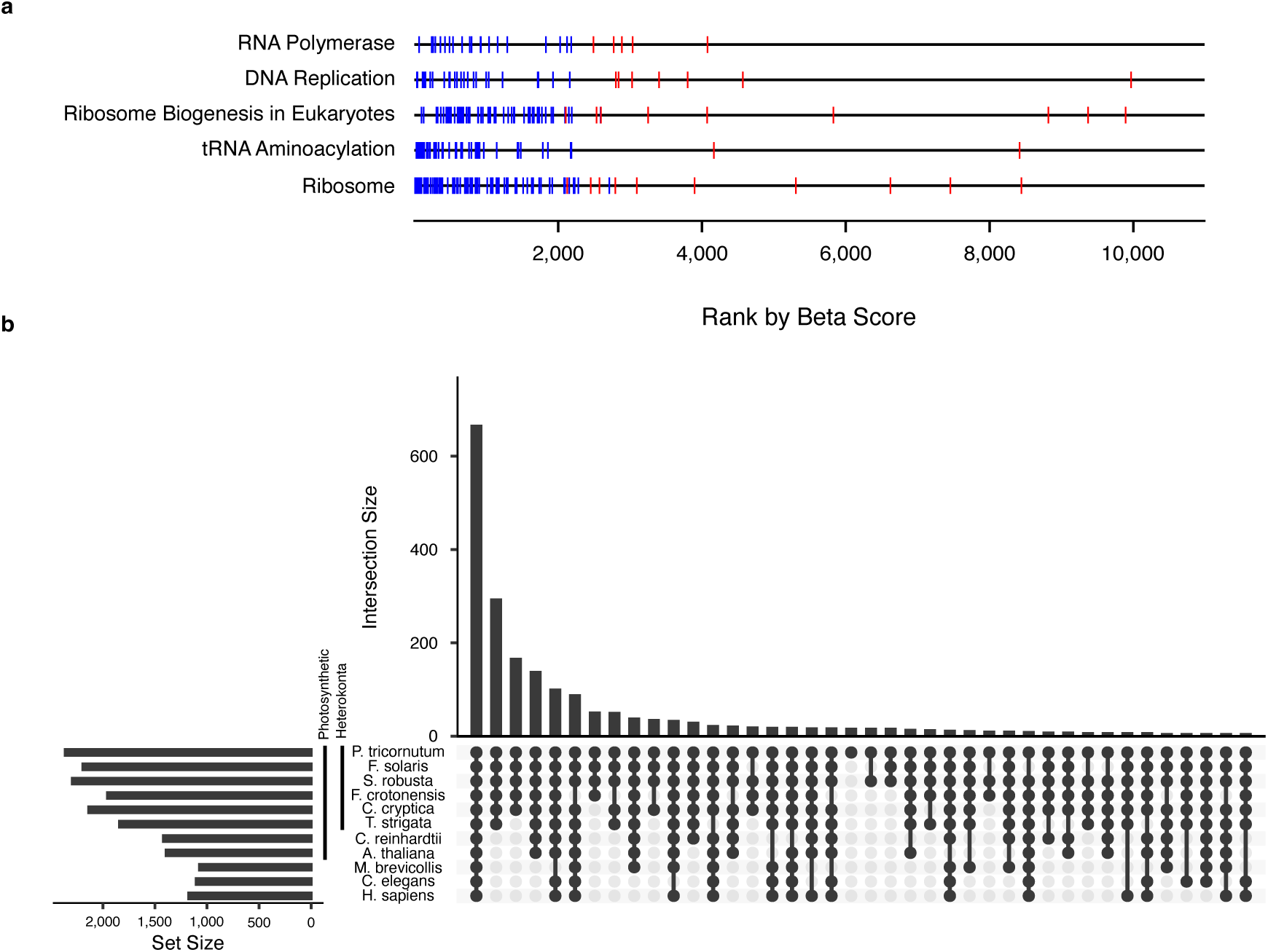
Characterization of *P. tricornutum* essential gene set. **a**, Recovery of genes from representative essential pathways. Genes are represented by vertical lines ranked according to their estimated gene essentiality (beta-score) and are classified as either essential (blue) or nonessential (red) in the screen. **b**, Conservation of *P. tricornutum* essential genes with other eukaryotes. Upset plot shows the number of *P. tricornutum* essential genes shared with specified eukaryotic groups (filled black circles).

**Extended Data Figure 4.**
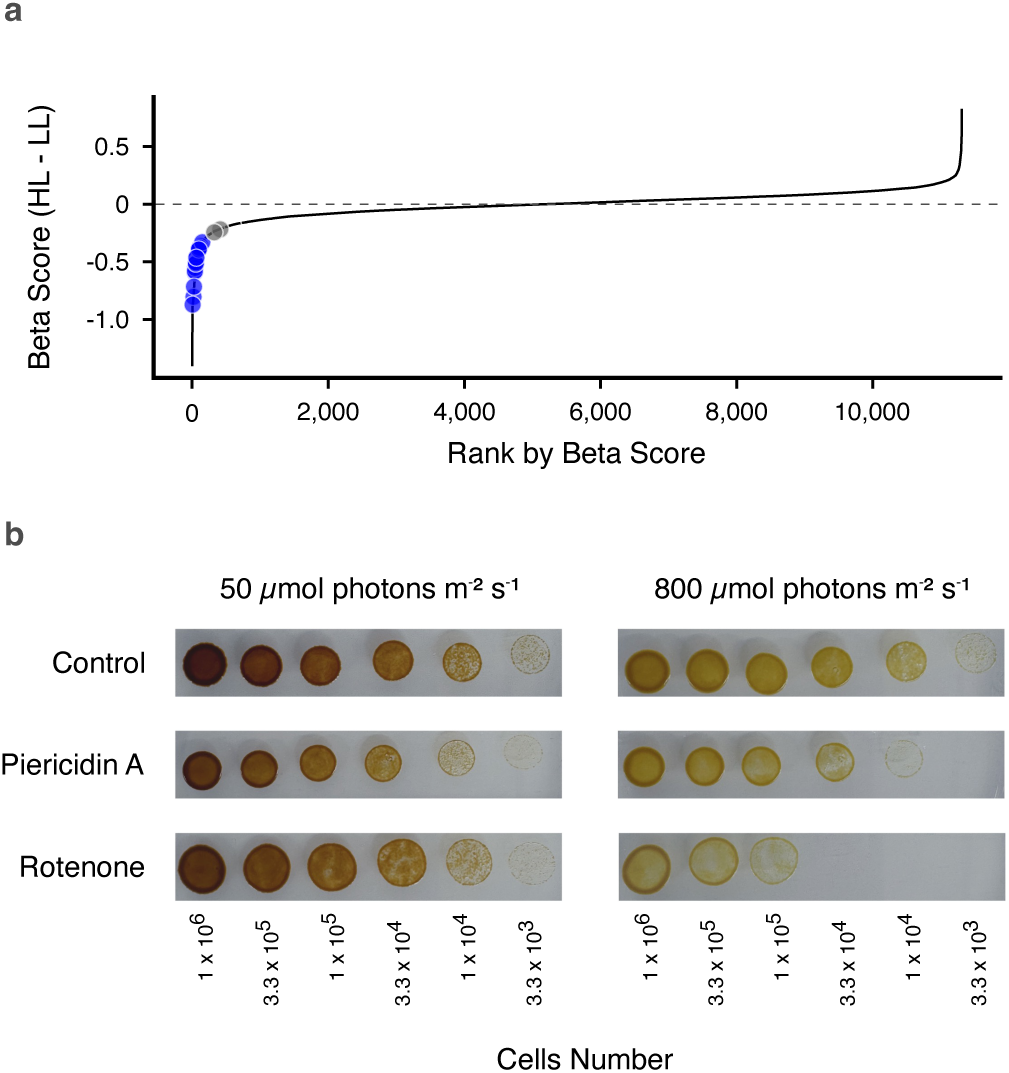
CMEF is required for survival under high light. **a**, Depletion of mitochondrial complex I subunits in high vs low light conditions. Gene rank (x-axis) is plotted against change in gene beta score (y-axis) in high light compared to low light conditions (black line). The mitochondrial complex I subunits encoded in the nuclear genome (circles) were identified in the high light depleted gene set (blue) or not identified in the high light depleted gene set (grey). **b**, Growth of *P. tricornutum* treated with mitochondrial Complex I inhibitors. Since the combined use of the respiratory inhibitors SHAM and Myxo has pleiotropic long-term effects, we used two independent mitochondrial complex I inhibitors, rotenone and piericidin. Cells were plated at different cell densities indicated and grown for 5 days in the following conditions: control untreated (top), piericidin A (middle, 5µM), or and rotenone (bottom, 10µM) under 50 µmol photons m^-2^ s^-1^ (left) or 800 µmol photons m^-2^ s^-1^ light (right). Images are representative of 2 biological replicates.

**Extended Data Figure 5.**
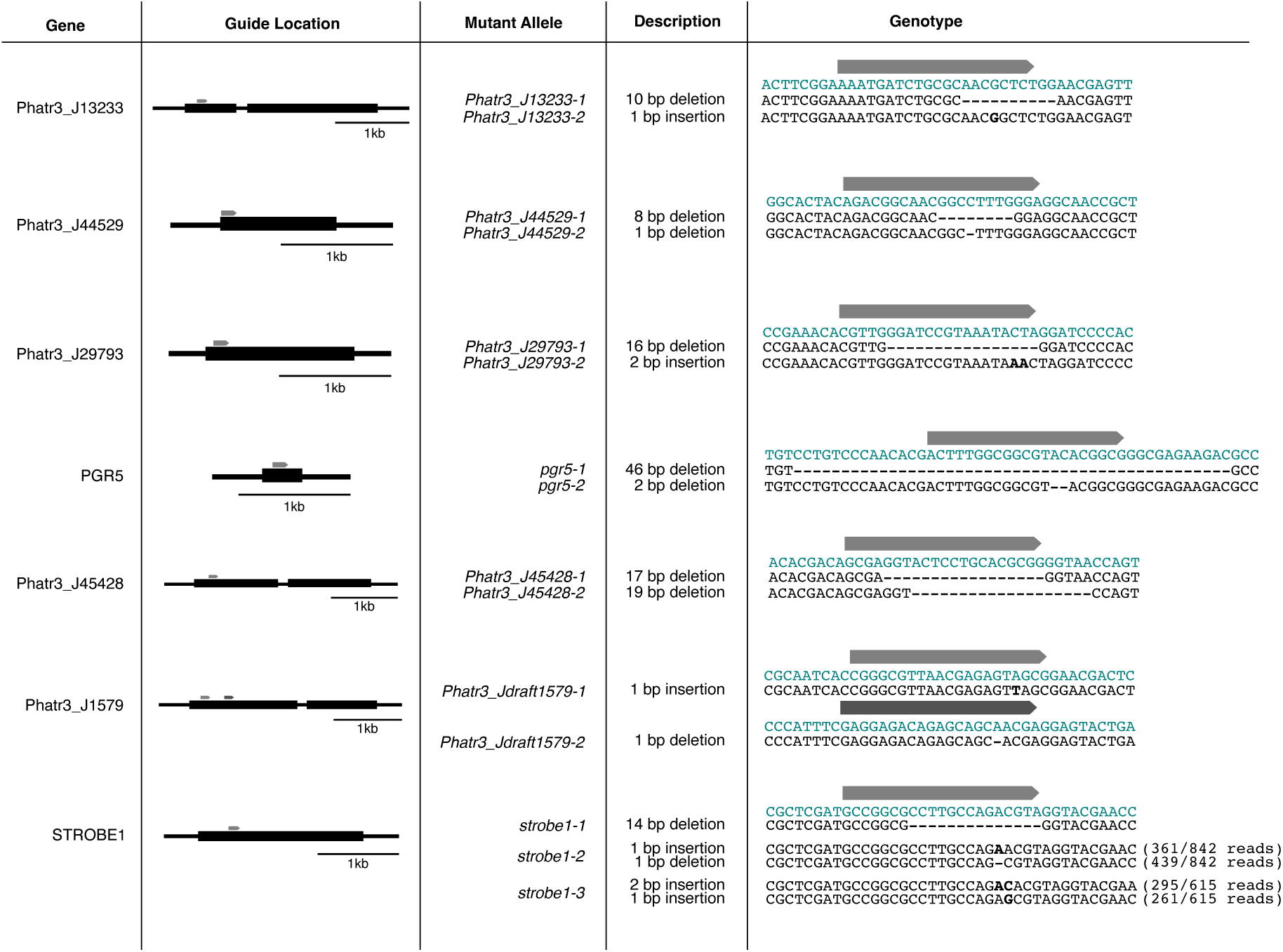
Disruption of dynamic light-specific genes. sgRNA design and mutation analysis for select genes depleted under dynamic light. Each row shows the targeted gene, location of the sgRNA (gray) in the annotated gene, mutant allele names, and resulting mutation type and mutant sequences for each genotype. Mutant sequences (black) are aligned to the reference sequence (teal). All genotypes were verified by Sanger sequencing; heterozygous edits were further verified by nanopore sequencing, with read counts reported for each allele.

**Extended Data Figure 6.**
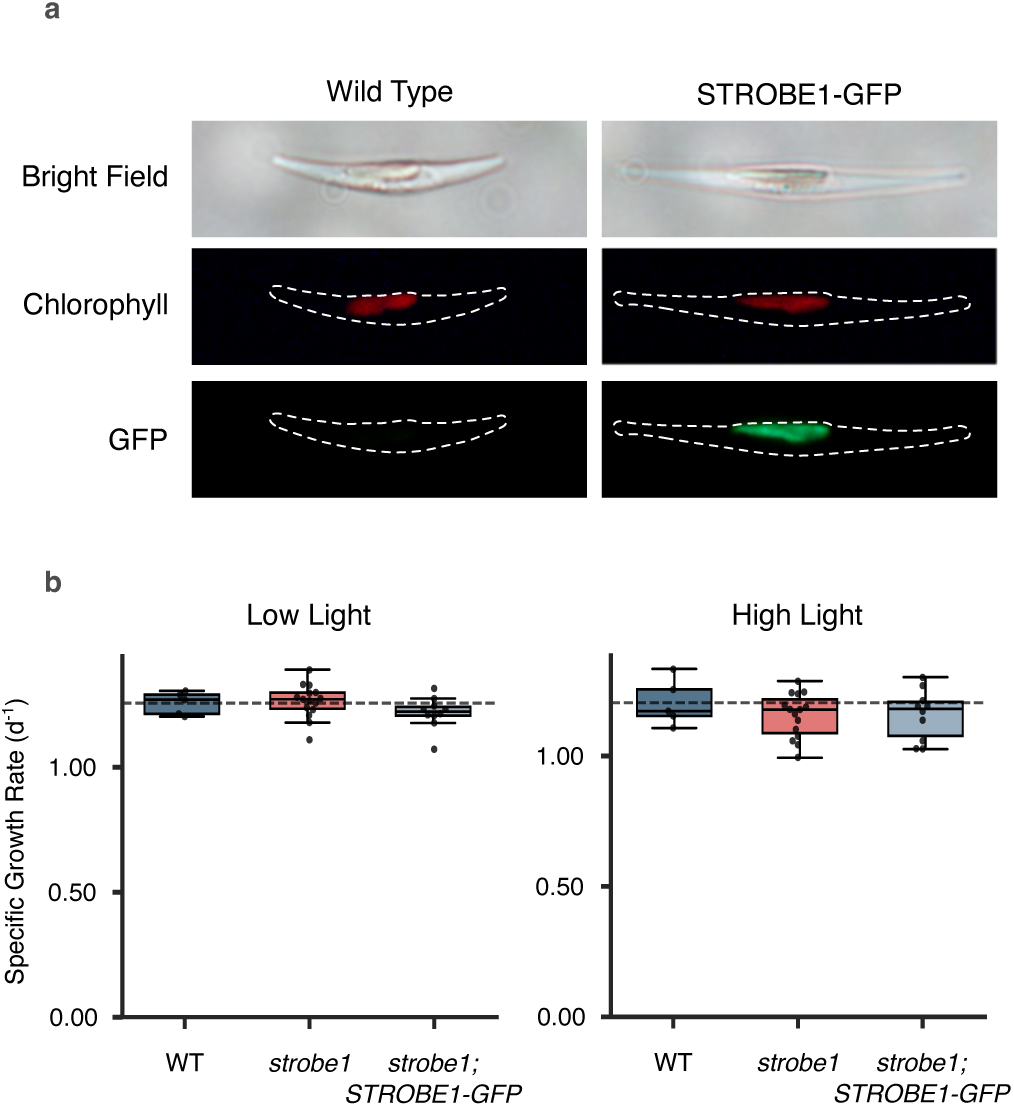
STROBE1-GFP localizes to the chloroplast and functionally complements *strobe1* mutants. **a**, Fluorescence images of wildtype (left panels) and *strobe1;STROBE1-GFP* (right panels) showing brightfield (top), chlorophyll autofluorescence (middle), and GFP (bottom). **b**, Specific growth rate of wildtype (dark blue), *strobe1* (red) and *strobe1*;*STROBE-GFP* (light blue) in low (50 µmol photons m^-2^ s^-1^) and high light (800 µmol photons m^-2^ s^-1^) growth conditions. Results shown are median (center line), interquartile-range (IQR, box) and 1.5xIQR (whiskers) for 5 biological replicates of ≥2 independent mutants. Dashed line represents mean from WT cells.

**Extended Data Figure 7.**
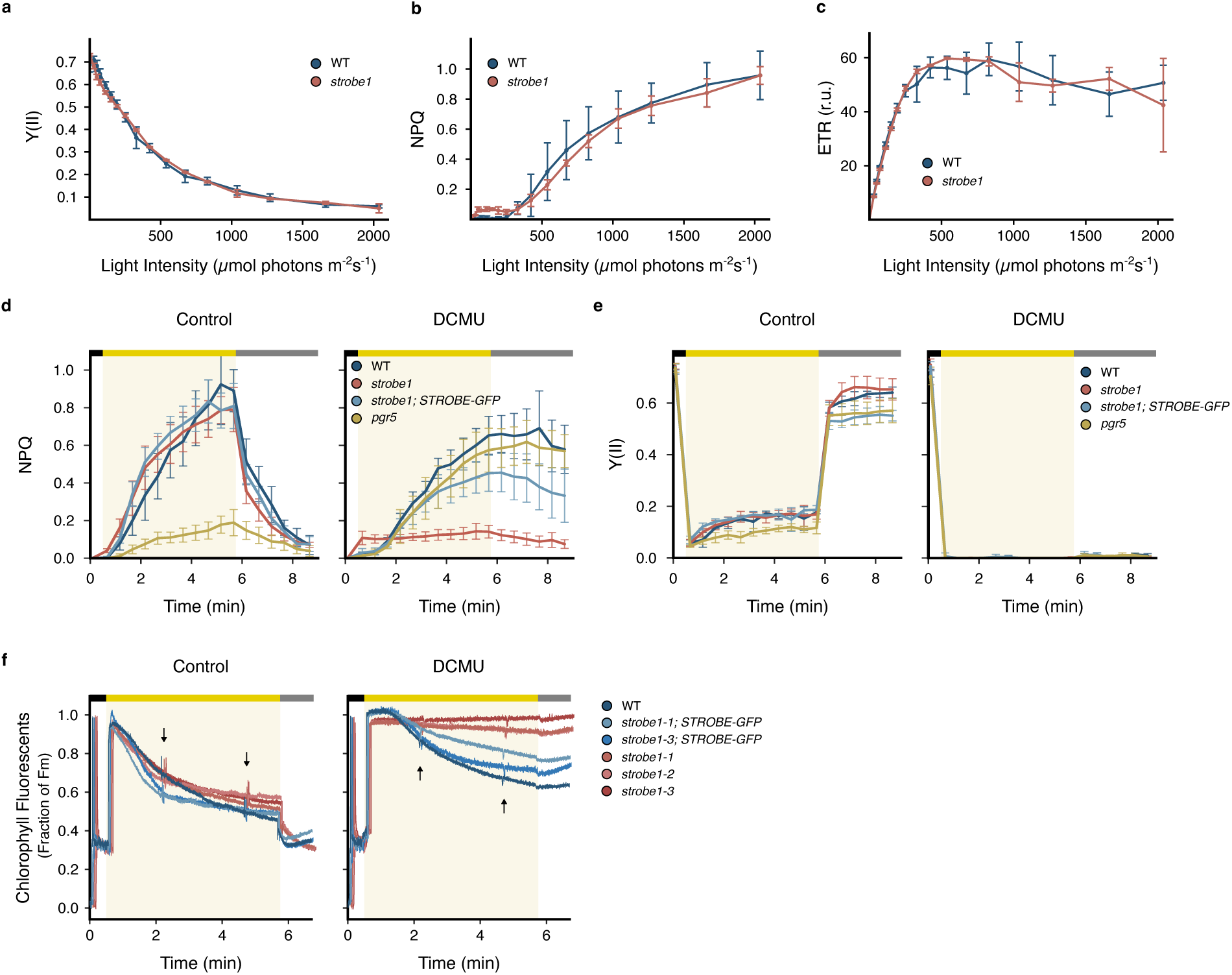
Chlorophyll fluorescence measurements for *strobe1* and *pgr5*. **a-c**, PSII yield (Y(II); **a**), NPQ (**b**), and relative PSII electron transport rate (ETR; **c**) after 1 minute of illumination at different light intensities. Results shown are mean +/- 1 standard deviation for 3 biological replicates for 3 independent mutants. **d-e,** NPQ (**d**) and PSII yield (**e**) measured during a dark to high light to low light transition. WT (dark blue), *strobe1* (red), *strobe1*;*STROBE-GFP* (light blue), and *pgr5* (yellow) were tested in absence (left panels) and presence of DCMU (10µM final concentration, right panels). Results shown are mean +/- 1 standard deviation for 3 biological replicates of ≥2 independent mutants. **f,** Typical chlorophyll fluorescence traces measured during a dark to high light to low light transition. Saturating pulses are shown with black arrows. WT (dark blue), independent *strobe1* mutants (shades of red), and independent *strobe1*;*STROBE-GFP* mutants (shades of blue) were tested in the absence (left panel) or presence of DCMU (10µM final concentration, right panel). Traces are staggered to avoid overlap. In **d-f**, yellow bars and shading indicate high light (800 µmol photons m^-2^ s^-1^), gray bars indicate low light (20 µmol photons m^-2^ s^-1^), and black bars indicate dark.

**Extended Data Figure 8.**
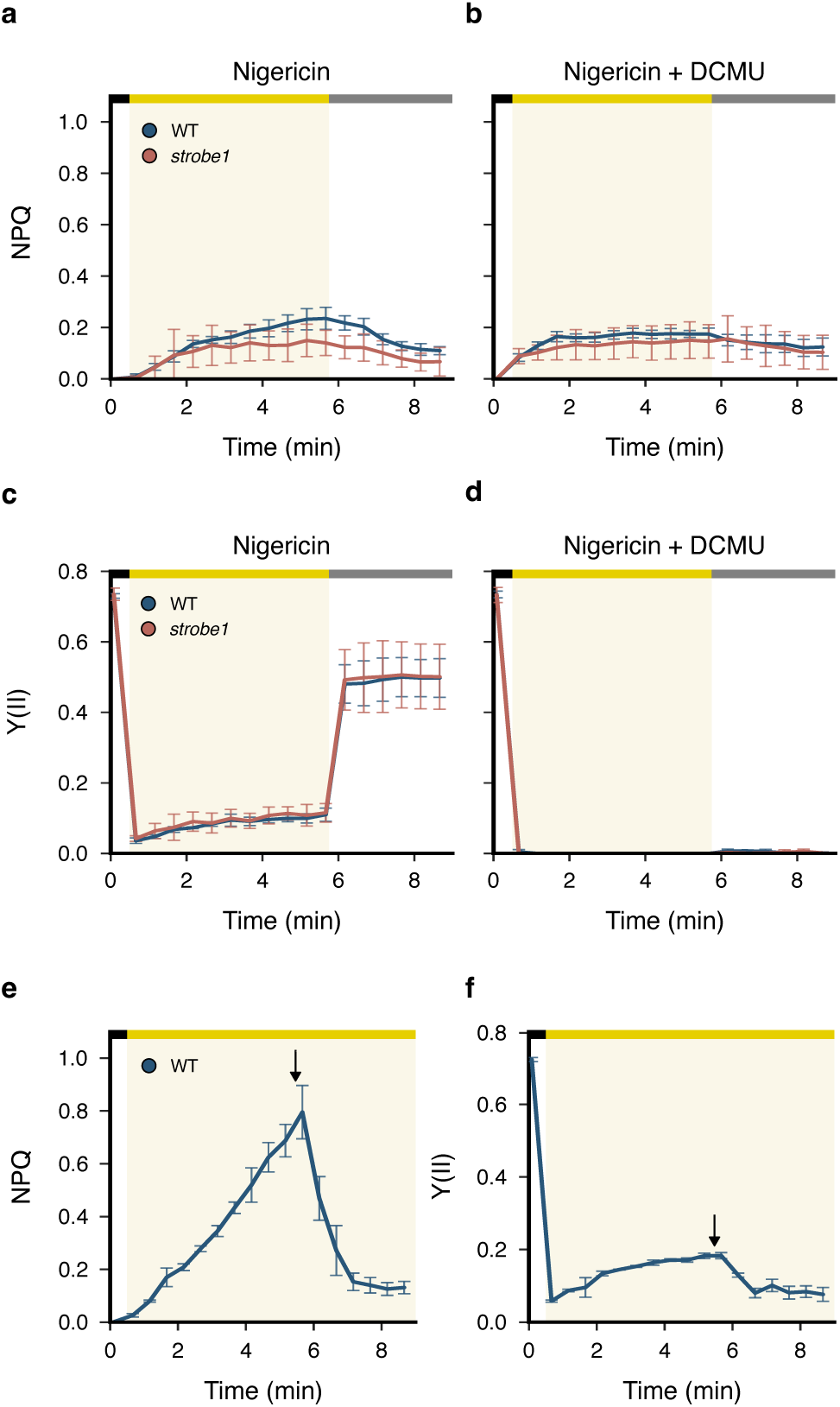
Nigericin dissipates the proton gradient and abolishes NPQ. **a-d.** NPQ (**a, b**) and PSII yield (Y(II); **c, d**) measured in the presence of the H^+^/K^+^ exchanger nigericin (100 µM final concentration), during a dark to high light to low light transition. Nigericin treated WT (dark blue) and *strobe1* (red) were tested in the absence (**a, c**) or presence (**b, d**) of DCMU (10µM final concentration). Results shown are mean +/- 1 standard deviation for 3 biological replicates of 3 independent mutants. **e-f** NPQ (**e**) and PSII yield (**f**) measured in WT cells during a dark to high light transition with addition of nigericin (100 µM final concentration) during the experiment as indicated by arrow. Results shown are mean +/- 1 standard deviation for 3 biological replicates. In **a-f**, yellow bars and shading indicate high light (800 µmol photons m^-2^ s^-1^), gray bars indicate low light (20 µmol photons m^-2^ s^-1^), and black bars indicate dark.

**Extended Data Figure 9.**
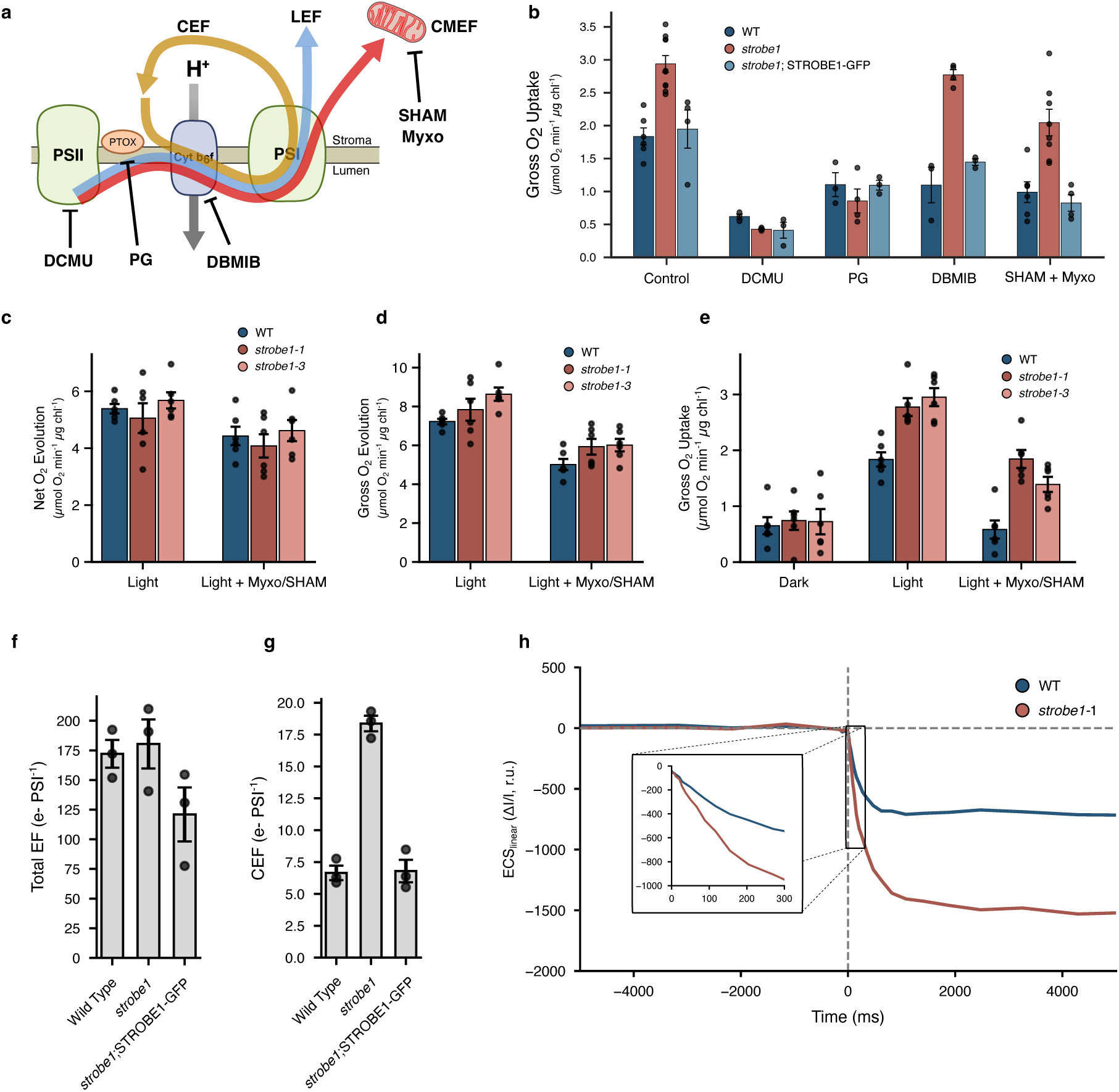
Characterization of photosynthetic electron flow in *strobe1* mutants using gas exchange measurements and electrochromatic shift. **a**, Diagram of the electron-transfer routes and inhibitors used: linear electron flow (LEF, blue); cyclic electron flow (CEF, orange); chloroplast–mitochondria electron flow (CMEF, red). The plastid terminal oxidase (PTOX, gray) can also facilitate additional electron flows. Inhibitors of these electron flows can be utilized to isolate electron flow mechanisms. PSII is inhibited with 3-(3,4-dichlorophenyl)-1,1-dimethylurea (DCMU), cytochrome (cyt) b_6_f is inhibited by 2,5-dibromo-3-methyl-6-isopropylbenzoquinone (DBMIB), mitochondrial respiration is inhibited by salicylhydroxamic acid (SHAM) and myxothiazol (Myxo), and PTOX is inhibited with propyl gallate (PG). H⁺ arrows indicate proton translocation driving the lumenal proton-dependent NPQ. **b-e**, Gas exchange rates were measured using a membrane-inlet mass spectrometry (MIMS) during a dark–to-light transition in wildtype (blue), *strobe1* (red) and *strobe1*;*STROBE1-GFP* (light blue) in the presence of absence of photosynthetic inhibitors. Gross O_2_ uptake rates (**b**) measured after 5 minutes of illumination when treated with indicated inhibitor. Results shown are ≥3 biological replicates for 2 independent *strobe1* mutant and one mutant of *strobe1*;*STROBE1-GFP*. Net O_2_ evolution (**c**), gross O_2_ evolution (**d**), and gross O_2_ uptake (**e**) for WT (blue) or 2 strains of *strobe-1* mutants (light and dark red) in the following conditions: dark, light, or light treated with respirations inhibitors SHAM and Myxo. Results shown are mean +/- 1 standard deviation of 6 biological replicates. **f-h**, Electron flow through the thylakoid membranes measured using electrochromic shift. WT, *strobe1*, and *strobe1*;*STROBE- GFP* were tested in the absence (**f**) or presence (**g**) of DCMU (10µM final concentration). In the absence of inhibitors, total electron flow (EF) is measured (**f**). DCMU treatment allows for the measurement of cyclic electron flow (CEF) (**g**). Results shown are mean +/- 1 standard deviation for 3 biological replicates of 1 *strobe1* strain and 1 *strobe1*;*STROBE1-GFP* strain. **h**, Representative ECS traces for WT (blue) and *strobe1* (red) upon a light to dark transition (t=0, vertical dashed line). Inset shows early kinetics of ECS immediately after light is turned off.

**Extended Data Figure 10.**
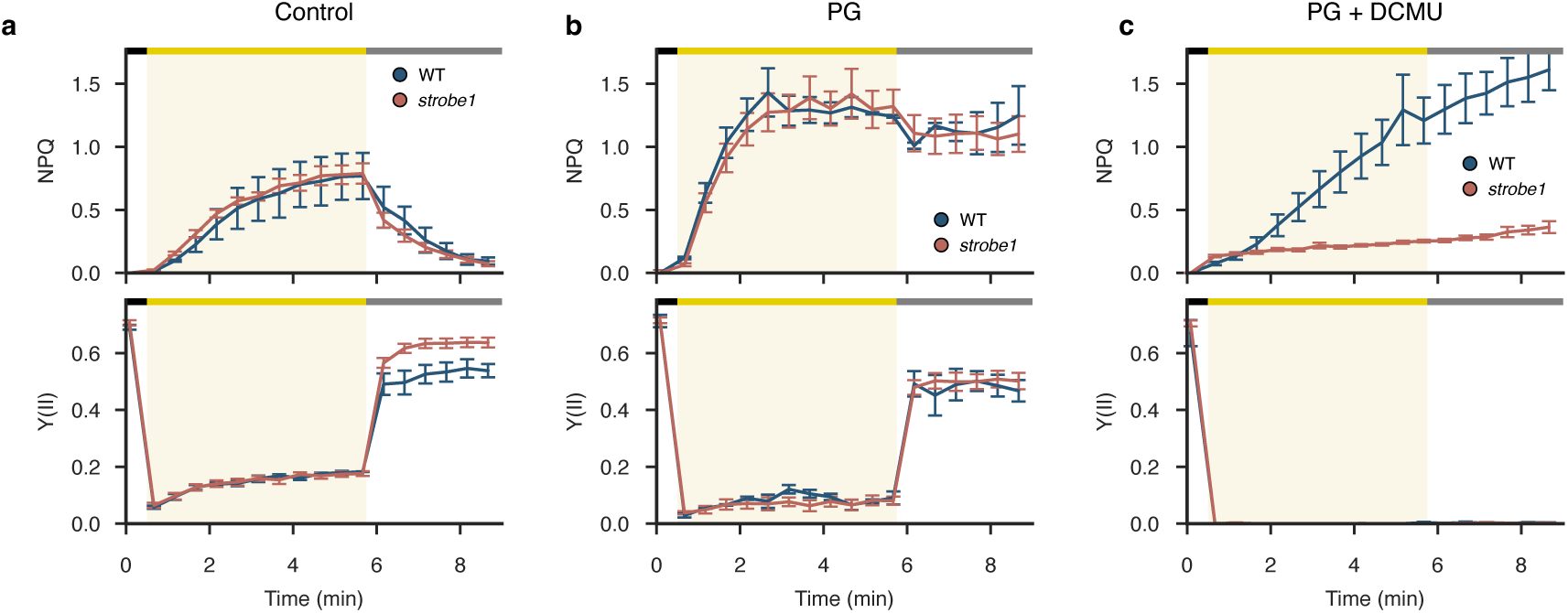
Chlorophyll fluorescence measurements for *strobe1* during PG inhibition with Propyl Galate. **a-c**, NPQ (top panels) and PSII yield (bottom panels). WT (blue) and *strobe1* (red) were tested in the following conditions: untreated control (**a**), PG (**b**), or PG + DCMU (**c**). Results shown are mean +/- 1 standard deviation for 3 biological replicates of 3 independent *strobe1* mutants. Yellow bars and yellow shading indicate high light (800 µmol photons m^-2^ s^-1^), gray bars indicate low light (20 µmol photons m^-2^ s^-1^), and black bars indicate dark.

**Supplementary Figure 1.**
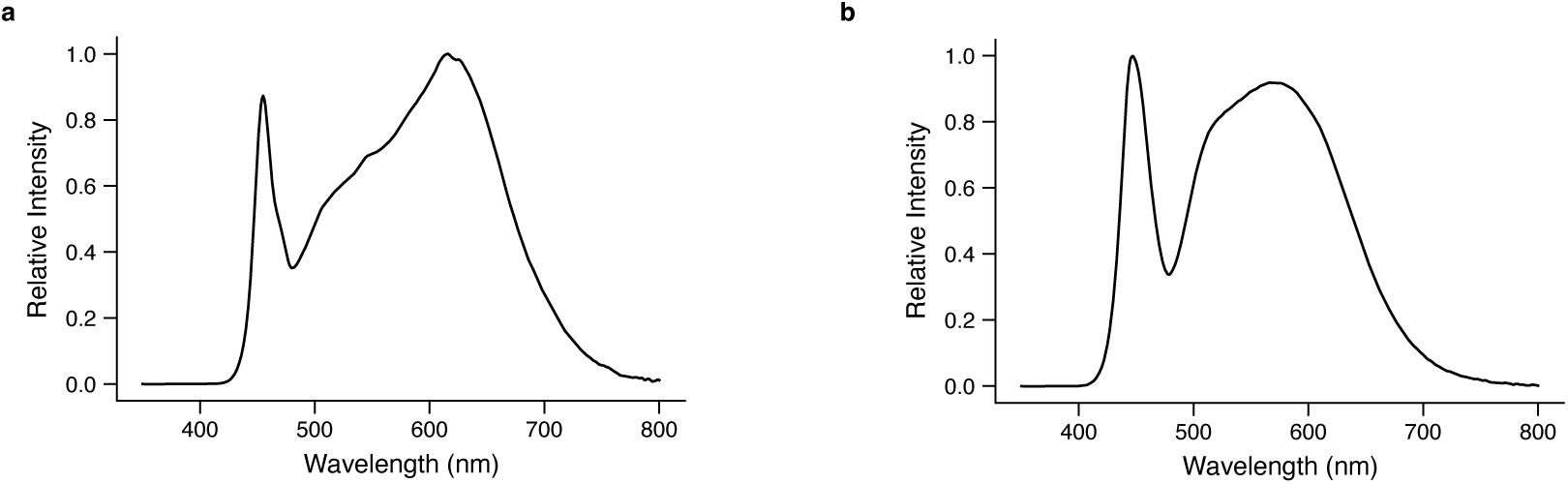
**a.** Light spectra of white LEDs used for all culturing and experiments of *P. tricornutum* unless otherwise stated. **b.** Light spectra of white LEDs used for genome-wide CRISPR/cas9 screen experiments and specific growth rate experiments.

**Supplementary Discussion**, including calculation of electron flows to support models of STROBE1 function

**Supplementary Table 1.** *Phaeodactylum tricornutum* sgRNA library sequences

**Supplementary Table 2.** *Phaeodactylum tricornutum* essential genes

**Supplementary Table 3.** *Phaeodactylum tricornutum* gene essentiality scores across all light treatments

**Supplementary Table 4.** High light-depleted genes in *Phaeodactylum tricornutum*

**Supplementary Table 5.** Dynamic light-specific genes in *Phaeodactylum tricornutum*

