## Supplemental Discussion for "A genome-wide CRISPR screen reveals how diatoms thrive in dynamic light"

### Supplementary Discussion

#### Quantification of the molecular mechanism by which STROBE1 manages lumenal proton

The different NPQ phenotypes of the *pgr5* and *strobe1* mutants when subjected to DCMU suggest that STROBE1's function might be independent of PGR5 and hence be an new potentiator of a protogenic CEF. However, this does not rule out the possibility of STROBE1 being involved in controlling a leak of the lumenal protons directly. We therefore aimed at developing a model to estimate the quantity of protons translocated by all photosynthetic electron flows in the WT and *strobe1* mutant. By knowing the rate of activity of each alternative photosynthetic electron flow, thanks to the capacity of each of them to translocate a proton, one can estimate the proton influx in the lumen and deduce what (i) the efficacy of a STROBE1-controlled CEF or (ii) the leak controlled by STROBE1 could be.

Using ECS and MIMS measurements, one can estimate the rate of each alternative photosynthetic electron flow relative to net photosynthesis (*%-LEF*)[1] at steady-state conditions. Under steady state conditions, net photosynthetic $O_2$ production is similar for the WT and *strobe1* mutant at 5 µmol $O_2$ min$^{-1}$ (Extended Data Fig. 9c). Comparison of gross $O_2$ evolution between control and treated samples with Myxo and SHAM (Extended Data Fig. 9d) shows CMEF to be around 1 µmol $O_2$ min$^{-1}$ µg Chl$^{-1}$ for both the WT and *strobe1* mutant, representing 20*%-LEF* (Supplementary Discussion Table 1 below). For PTOX activity, comparison of gross $O_2$ uptake between control and treated samples with propyl gallate (PG, Extended Data Fig. 10) shows PTOX activity to be around 1 µmol $O_2$ min$^{-1}$ µg Chl$^{-1}$ for the WT and 2 µmol $O_2$ min$^{-1}$ µg Chl$^{-1}$ for the *strobe1* mutant, representing 20 and 40*%-LEF,* respectively (Supplementary Discussion Table 1 below). As addition of the PTOX inhibitor PG did not change the strobe1 mutant's fluorescence phenotypes (Extended Data Fig. 10), the increased PTOX activity in *strobe1* mutants is not causative of the *strobe1* phenotype and is likely due to higher CEF occurring in the *strobe1* mutant which creates a stronger reduction of the plastoquinone pool. CEF measurements in the presence of DCMU show a rate of 6 and 18 e$^-$ s$^{-1}$ PSI$^{-1}$ for the WT and *strobe1* mutant, respectively (Extended Data Fig. 9h). Note that those measurements reflect maximal CEF activity, and not necessarily CEF rates in physiological conditions. However, a higher CEF rate in *strobe1* as compared to the WT is compatible with a higher PTOX activity; one could thus estimate that CEF in physiological conditions in the *strobe1* mutant is < 18 e$^-$ s$^{-1}$ PS$^{-1}$ and > 6 e$^-$ s$^{-1}$ PS$^{-1}$. To estimate those rates in *%-LEF*, we used ECS to estimate total electron flow (TEF) in the absence of DCMU (**Extended Data Fig. 9g-i**), being respectively 124 and 144 e$^-$ s$^{-1}$ PS$^{-1}$. Since LEF and CMEF translocate 1e$^-$ PS$^{-1}$, PTOX translocates 1e$^-$ PSII$^{-1}$, CEF translocates 1e$^-$ PSI$^{-1}$; thus TEF relates to photosynthetic electron flows as:

$$(1)\ TEF=LEF+CMEF+1/2\times PTOX+1/2\times CEF$$

Using in (1) the percentage of each electron flow in *%-LEF* and the values of CEF measured in the presence of DCMU, we estimate LEF to be ~ 90 $e^{-}$ $s^{-1}$ $PS^{-1}$, in the WT and *strobe1*. CEF might therefore represent 7*%-LEF* in the WT and between 7-17*%-LEF* in the *strobe1* mutant.
The steady-state proton translocation rate mediated by photosynthetic electron flow in each strain can then be computed as:

$$(2)\ H^{+}_{in}=LEF\times(Y_{LEF}+CMEF_{\%\text{-}LEF}\times Y_{CMEF}/100+PTOX_{\%\text{-}LEF}\times Y_{PTOX}/100+CEF_{\%\text{-}LEF}\times Y_{CEF}/100)$$

Where $EF_{\%\text{-}LEF}$ is the electron flow expressed in *%-LEF* and $Y_{EF}$ is the capacity of each electron flow to create a lumenal $H^+$ per electron[2]. We can now use (2) to test the boundaries of each hypothesis, to see if (i) a STROBE1-mediated CEF would be required to pump a reasonable amount of $H^+$ per electron or (ii) a STROBE1-controled $H^+$ leak would be required to regulate a reasonable flow of $H^+$.

**Hypothesis I: STROBE1 mediates a CEF.** In this case, the context which would require STROBE1-mediated CEF to pump the least amount of $H^+$ per electron would be if the STROBE1-mediated CEF is the main CEF active in the WT while only the PGR5-controlled one would be active in the *strobe1* mutant. We assume here that the leak of protons out of the thylakoid lumen and that the accumulation of lumenal protons is similar in the WT and the *strobe1* mutant (Fig. 3). Under this hypothesis, the *strobe1* mutant harbors a $H^{+}_{in}$ of ~372-390 $H^+$ $s^{-1}$ $PS^{-1}$ for the low and high CEF scenario respectively, while the WT without CEF shows a $H^{+}_{in}$ of 342 $H^+$ $s^{-1}$ $PS^{-1}$. Hence, 7*%-LEF* of STROBE1-mediated CEF activity measured in the WT should allow compensating for the difference of $H^{+}_{in}$ between the WT without CEF and *strobe1*, ( 30-41 $H^+$ $s^{-1}$ $PS^{-1}$), which, using (2), corresponds to a STROBE1-mediated CEF capacity to create a lumenal $H^+$ per electron ($Y_{STROBE1}$) between 4.7 and 6.5 $H^+$ per electron.
Note that uncertainties in the estimate of CEF in physiological conditions might affect these values; however, the scenario that would give us the lower $Y_{STROBE1}$ would be if CEF in *strobe1* was zero and CEF in the WT was maximal and mediated only by STROBE1. Although unrealistic, this would still lead to a $Y_{STROBE1}$ of 2.8 $H^+$ per electron, a value higher than the 2 $H^+$ per electron mediated by the PGR5-controlled CEF.

**Hypothesis II: STROBE1 controls a leak of protons through the thylakoid membrane.** In this case, we hypothesize that the main CEF active in the WT and the *strobe1* mutant is PGR5-controlled. We assume here that the accumulation of lumenal protons is similar in the WT and the *strobe1* mutant (Fig. 3). Under this hypothesis, the *strobe1* mutant still harbors a $H^{+}_{in}$ of 372-390 $H^+$ $s^{-1}$ $PS^{-1}$, while the WT shows a $H^{+}_{in}$ of 354 $H^+$ $s^{-1}$ $PS^{-1}$, which corresponds to STROBE1 controlling a leak of 18-36 $H^+$ $s^{-1}$ $PS^{-1}$, corresponding to a controlled leak of 5-10% of the pumped protons.

These calculations reveal that if STROBE1 mediates a CEF, it should produce more protons per electron than the PRG5-controlled CEF. The NDH-mediated CEF is the most efficient CEF known, with a $Y_{NDH}$ of 4 $H^+$ per electron [3] the $Y_{STROBE1}$ estimated here is on the high end of this value, suggesting that STROBE1 would be part of a larger complex that can highly efficiently translocate protons. On the other hand, a proton leak of 5-10% could be in line with a potential presence of (i) one or two uncoupling proteins per PS, like the mitochondrial UCP1, for which, when isolated from Syrian hamsters, transport 14 $H^+$ $s^{-1}$ per transporter[4] or (ii) a modulation of the activity of the ATP synthase[5]. Regardless of how STROBE1 regulates the lumenal proton concentration, its presence is important to quickly build a low lumenal pH in a CEF-dependent manner, leading to increased photoprotection and ATP synthesis in dynamic light environments.

| Electron flow | Thylakoid lumenal $H^+$ created per electron | Activity in the WT (*%-LEF*) | Activity in *strobe1* (*%-LEF*) |
|---|---|---|---|
| LEF | 3 | 100 | 100 |
| CMEF | 3 | 20 | 20 |
| PTOX | 1 | 20 | 40 |
| CEF | 2 (PGR5-controlled) | 7 | 7-17 |

**Supplementary Discussion Table 1. Activity of each electron flow in the WT and *strobe1* mutant and their capacity to generate lumenal $H^+$.** $H^+$ per electron coupling efficiencies of photosynthetic electron flow are drawn based on previously estimates[2,3]. The activity of each photosynthetic electron flow is estimated in the text above from Extended Data Fig. 9 and 10.
